# The order Axinellida (Porifera: Demospongiae) in California

**DOI:** 10.1101/2022.03.02.482672

**Authors:** Thomas L. Turner, M. Sabrina Pankey

## Abstract

Sponges are common and diverse in California, but they have received little study in the region, and the identities of many common species remain unclear. Here we combine fresh collections and museum vouchers to revise the order Axinellida for California. Seven new species are described: *Endectyon (Endectyon) hispitumulus, Eurypon curvoclavus, Aulospongus viridans, Aulospongus lajollaensis, Halicnemia litorea*, *Halicnemia montereyensis*, and *Halicnemia weltoni*. One new combination is also described, and two existing species are reduced to junior synonyms, resulting in a total of 13 species; a dichotomous key to differentiate them is provided. DNA data from 9 of the 13 species is combined with publicly available data to produce updated global phylogenies for the order.

## Introduction

The order Axinellida (Levi, 1953) is a large group of demosponges with a world-wide distribution. In its current iteration, the order is comprised of four families: the Axinellidae (Carter 1875), Raspailiidae (Nardo 1833), Stelligeridae (von Lendenfeld 1898), and Heteroxyidae (Dendy 1905) (Morrow *et al*. 2019; Morrow & Cárdenas 2015), containing an estimated 669 described species across 49 genera (de Voogd *et al*. 2022). The order was originally proposed by Lévi (1953), who hypothesized a close relationship between the families Axinellidae and Raspailiidae, but the order underwent numerous revisions and was eventually disbanded due to lack of supporting morphological evidence (Alvarez & Hooper 2002; van Soest *et al*. 1990). DNA phylogenies later verified the close relationship between Axinellidae and Raspailiidae, and the order was resurrected (Morrow & Cárdenas 2015). Genetic data further led to the restoration of the family Stelligeridae and the inclusion of that family, and the Heteroxyidae, in the Axinellida (Morrow *et al*. 2012). Understanding the relationships among these four families and their component genera is a work in progress, as molecular phylogenies indicate that many taxa are polyphyletic (Carballo *et al*. 2018; Gazave *et al*. 2010; Redmond *et al*. 2013; Thacker *et al*. 2013). The recent history of sponge taxonomy makes it clear that DNA data will be a major component of continued revisions, but this presents a challenge, as many sponges in museum collections have failed to yield amplifiable DNA (Vargas *et al*. 2012; also see methods below). It is therefore likely that fresh collections, which are both sequenced and morphologically described, will be crucial to continued improvement.

Six species of Axinellida were previously known from California (Lee *et al*. 2007a). Half were named by de Laubenfels in the 1930s (de Laubenfels 1932, 1935), while the others were described by Dickinson (Dickinson 1945) and Sim and Bakus (Sim & Bakus 1986). Most were described from only one or two samples. Additional samples from the Mexican Pacific have increased our understanding of some of these species (Aguilar-Camacho & Carballo 2013; Gómez *et al*. 2002), but we still know little about their status and distribution, and DNA data is absent for most. In addition to the gaps in our knowledge of described species, other morphologies have been discovered but not described (Lee *et al*. 2007b; a), while an unknown number also await discovery.

To fill these gaps in our knowledge, we have led an effort to collect and document the sponges of California, largely via SCUBA diving. To date, this ongoing project has netted 1155 sponge samples, mostly from diving depths but also from the intertidal zone and marinas. We previously used this collection to revise the orders Tethyida (Turner 2020) and Scopalinida (Turner 2021), and have now identified 73 of these samples as members of the Axinellida. These include fresh collections of all but one of the named shallow-water species, an additional species that was known to exist but had not been described, and three additional species that were previously unknown. By analyzing this collection together with 20 samples from existing museum collections, we are able to present a comprehensive look at the California Axinellida, including in situ photos of live sponges, qualitative data on abundance and ranges, and quantitative spicule data. DNA sequence from 9 of the species is analyzed together with publicly available genetic data to produce updated phylogenies for the order.

## Methods

### Collections

Sponge samples were collected by hand with a knife (*Dragmacidon mexicanum* (n=32), *Endectyon (Endectyon) hyle* comb. nov. (n=9), *Endectyon (Endectyon) hispitumulus* sp. nov. (n=9), *Eurypon curvoclavus* sp. nov. (n=3), *Aulospongus viridans* sp. nov. (n=1), *Cyamon neon* (n=11), *Trikentrion helium* (n=2), *Halicnemia litorea* sp. nov. (n=4)). Samples were placed in one-liter plastic bags with seawater, and these were kept on ice until sponges could be preserved in 95% EtOH. Samples were given a collection ID (numbers beginning with TLT) until they could be vouchered in a permanent collection. Holotypes and some other samples were vouchered with the California Academy of Sciences in San Francisco (voucher numbers with CASIZ). Additional samples were vouchered with the Santa Barbara Museum of Natural History (voucher numbers with SBMNH) and the Cheadle Center at the University of California, Santa Barbara (voucher numbers with UCSB). Several samples were collected during the “Los Angeles Urban Ocean Expedition 2019” bioblitz; these are archived at both the Natural History Museum of Los Angeles (voucher numbers with NHMLA) and the Florida Museum of Natural History (voucher numbers with BULA). Additional samples were acquired on loan from the California Academy of Sciences, the Natural History Museum of Los Angeles County, the Santa Barbara Museum of Natural History, and the Scripps Institution of Oceanography (vouchers numbers beginning with P). Voucher and/or collection numbers are listed for each species in the systematic section, but we also provide them in tabular format as supplementary data deposited at Data Dryad (https://doi.org/10.25349/D93W5B). This supplementary table includes additional metadata for each sample, such as collection locations and Genbank numbers.

Seventy-nine subtidal locations were investigated while SCUBA diving: 14 around Monterey Bay and Carmel Bay, in Central California, and the rest in the Southern California Bight. Most sites (75%) were searched for only a single dive, but some areas, especially in the Santa Barbara Channel, were searched repeatedly. Sites varied from 6 to 33 m in depth; most (75%) were less than 22 m deep. Intertidal (14 sites) and harbor (23 sites) collections were less extensive, but covered a broader latitudinal range, from Drake’s Estero (Marin County, Northern California) to San Diego Bay, near the Southern border of California.

Collections at each location were focused on novel-looking sponges and sponges that cannot be identified in the field, while also photographically documenting all field-identifiable sponges present. This allows sponge presence/absence across sites to be compiled, and these data can be used to form hypotheses about sponge distributions and habitat. Field-identifiability of sponges was tested by attempting to identify each sponge using field characters, and then testing field identifications using spicules and/or DNA sequencing. The reliabilities of field-identifications are variable, based on the number of individuals tested and the consistency of their morphology, as indicated in the systematic section. Because search effort varied among sites, and a uniform protocol was not used, distribution data should be considered preliminary and non-quantitative. A supplementary table deposited at Data Dryad (https://doi.org/10.25349/D93W5B) contains additional information about locations investigated, such as an estimate of search effort per location and data on sponge presence/absence for each species; a map of collection locations is also provided.

### Morphology

Individual spicules were examined by digesting sponge samples in bleach. Skeletal architecture was investigated by hand cutting tissue sections and digesting them with a mixture of 97% Nuclei Lysis Solution (Promega; from the Wizard DNA isolation kit) and 3% 20mg/ml Proteinase K (Promega). This digestion eliminates cellular material while leaving the spongin network intact. Spicules and sections were imaged with a D3500 SLR camera (Nikon) with a NDPL-1 microscope adaptor (Amscope) attached to a compound trinocular microscope. Measurements were made on images using ImageJ (Schneider *et al*. 2012) after calculating the number of pixels per mm with a calibration slide. Unless otherwise specified, spicule length was measured as the longest possible straight line from tip to tip, even when spicules were curved or bent. Spicule width was measured at the widest point, excluding adornments like swollen tyles. Secondary electron images were taken with a FEI Quanta400F Mk2 after coating spicules with 20 nm of carbon. Scale bars on figures are precise for microscope images but often approximate for field photos. All spicule measurements are available as raw data at Data Dryad (https://doi.org/10.25349/D93W5B).

### Genotyping

DNA was extracted with several different kits: the Wizard Genomic DNA Purification kit (Promega), the Qiagen Blood & Tissue kit, and the Qiagen Powersoil kit. Though we have found that the Wizard kit works well on sponges from other sponge orders, PCR amplifications from these extractions usually failed from the sponges included here, even when extractions were diluted. Qiagen kits, which include a column filtration, were more successful, though extracts from the Blood & Tissue kit often required dilution. These differences are presumably due to co-purification of PCR inhibiting compounds in these species.

Two primer sets were used to sequence a fragment of the *cox1* mitochondrial locus. Most samples were genotyped at a ∼1200 bp fragment using the following primers (LCO1490: 5-GGT CAA CAA ATC ATA AAG AYA TYG G-3’; COX1-R1: 5’-TGT TGR GGG AAA AAR GTTAAA TT-3’); these amplify the “Folmer” barcoding region and the “co1-ext” region used by some sponge barcoding projects (Rot *et al*. 2006). The Folmer region alone was amplified from some samples using the following primers: (LCO1490: 5’-GGT CAA CAA ATC ATA AAG AYA TYG G-3’; HCO2198: 5’-TAA ACT TCA GGG TGA CCA AAR AAY CA-3’) (Folmer *et al*. 1994). Though both primer sets worked well for the Raspailliidae, neither worked in the Stelligeridae, and only one of two species of Axinellidae was successful. In fresh samples, the Folmer primers often produced an amplicon, but sequencing revealed it to be a mixed sample, presumably contaminated by the unicellular (or multicellular) symbionts abundant in sponges.

Three primer sets were used to amplify portions of the 28S rDNA nuclear locus. Most species were sequenced across the D1-D5 region by combining two amplicons: the ∼800 bp D1-D2 region using primers Por28S-15F (5’-GCG AGA TCA CCY GCT GAA T-3’) and Por28S-878R (5’-CAC TCC TTG GTC CGT GTT TC-3’) and the ∼800 bp D3-D5 region using primers Por28S-830F (5’-CAT CCG ACC CGT CTT GAA-3’) and Por28S-1520R (5’-GCT AGT TGATTC GGC AGG TG-3’) (Morrow *et al*. 2012). The D1-D2 primers failed to yield products for both *Endectyon* species. For these samples, the C2-D2 region (covering ∼50% of the D1-D2 region) was amplified with C2 (5’-GAA AAG AAC TTT GRA RAG AGA GT-3’) and D2 (5’-TCC GTG TTT CAA GAC GGG-3’) (Chombard *et al*. 1998).

PCR was performed using a Biorad thermocycler (T100); the following conditions were used for the cox1 locus: 95°C for 3 min, followed by 35 cycles of 94°C for 30 sec, 52°C for 30 sec, 72°C for 90 seconds, followed by 72°C for 5 minutes. The 28S C2-D2 region was amplified with the same conditions, except a 57°C annealing temperature and 60 second extension time; the 28S D1-D2 and D3-D5 regions used a 53°C annealing temperature and 60 second extension time. PCR was performed in 50 μl reactions using the following recipe: 24 μl nuclease-free water, 10 μl 5x PCR buffer (Gotaq flexi, Promega), 8 μl MgCl, 1 μl 10mM dNTPs (Promega), 2.5 μl of each primer at 10 μM, 0.75 bovine serum albumin (10 mg/ml, final conc 0.15 mg/ml), 0.25 μl Taq (Gotaq flexi, Promega), 1 μl template. ExoSAP-IT (Applied Biosystems) was used to clean PCRs, which were then sequenced by Functional Biosciences using Big Dye V3.1 on ABI 3730xl instruments. Blastn was used to verify that the resulting traces were of sponge origin. All sequences have been deposited in GenBank; accession numbers are shown in the phylogenies and also listed in the supplementary table in Data Dryad (https://doi.org/10.25349/D93W5B).

PCR was attempted from a variety of older museum vouchers and was nearly always unsuccessful. These include samples of multiple *Halicnemia* species from the California Academy of Sciences, *Raspailia hymani* from the Santa Barbara Museum of Natural History, and the holotypes of *Trikentrion helium*, *Cyamon argon*, *Dragmacidon oxeon*, and *Stylissa letra* from the Natural History Museum of Los Angeles. Most DNA extractions produced little detectable DNA, and PCR amplification of these samples failed to yield any bands at either locus. When PCR on these samples yielded a product, it was usually due to contamination, and this contamination appeared to be from past handling of the vouchers rather than contamination during molecular work. For example, the *C. argon* holotype produced a 28S sequence from a Tetractinellid, and the molecular lab performing PCR had not previously amplified sponge DNA from this order. Extractions performed on multiple *Halicnemia* from the California Academy of Sciences all produced a 28S sequence matching a common California *Suberites*. This could have result from contamination of the samples during molecular work, but these samples were not extracted as a set, and all were contaminated with the same sequence, so contamination of the vouchers by a past taxonomist seems most likely. The only museum sample that yielded DNA sequence that was likely native was a sample of *Halicnemia weltoni sp. nov.*. This sequence is both unique and has a phylogenetic position supported by its morphology.

### Phylogenetic methods

Axinellid sequences were collected from GenBank using a combination of blast and the NCBI taxonomy browser; effort was made to find every available sequence at these loci from sponges in the Axinellida (at 28S) or the Raspailiidae (at cox1), as well as relevant outgroups. For cox1, sequences were included if they minimally encompassed the Folmer barcoding region. For 28S, sequences were included if they minimally encompassed the C2-D2 barcoding region. Combined, the previously published sequences included in the phylogenies are from 19 different publications and several unpublished datasets (Belinky *et al*. 2012; Erpenbeck *et al*. 2007a; b, 2012, 2016; Gazave *et al*. 2010; Holmes & Blanch 2007; Huchon *et al*. 2015; Idan *et al*. 2018; Kayal & Lavrov 2008; Lavrov *et al*. 2008; Lim *et al*. 2017; Morrow *et al*. 2012, 2013, 2019; Nichols 2005; Richelle-Maurer *et al*. 2006; Schuster *et al*. 2021; Turon *et al*. 2018). *Hymeraphia vaceleti* 28S sequence was provided by Christine Morrow.

Sequence alignments were produced using MAFFT v.7 online (Katoh *et al*. 2017). Phylogenies were estimated with maximum likelihood using a GTR model in IQ-Tree (Nguyen *et al*. 2015; Trifinopoulos *et al*. 2016); the ultrafast bootstrap was used to measure node confidence (Hoang *et al*. 2018). Figures were produced by exporting IQ-Tree files to the Interactive Tree of Life webserver (Letunic & Bork 2019).

## Results

### Phylogenies

A phylogeny for the 28S ribosomal RNA locus is shown in figure 1. We successfully sequenced portions of this locus from 21 California samples, representing 9 of the 13 species of California Axinellida. A minimum of 1224 bp was sequenced from at least one sample from each species. We compare these new sequences to 91 publicly available sequences; these samples vary greatly in length (mean = 1626 bp), but were only included if they encompassed the highly variable C2-D2 region (∼ 500 bp) targeted by the Sponge Barcoding Project (Chombard *et al*. 1998). Our primary aim was to use these phylogenies as part of an integrative revision of California species, so we chose breadth (including as many species as possible) over depth (including only the species with the most data, to better resolve branching order). Though deeper branches in the phylogeny are poorly resolved, we were able to generate strong phylogenetic hypotheses for many important clades and confidently identify phylogenetic species. Note that the family Heteroxyidae is absent because no species in this family have data available at the 28S C2-D2 region.

**Figure 1.**
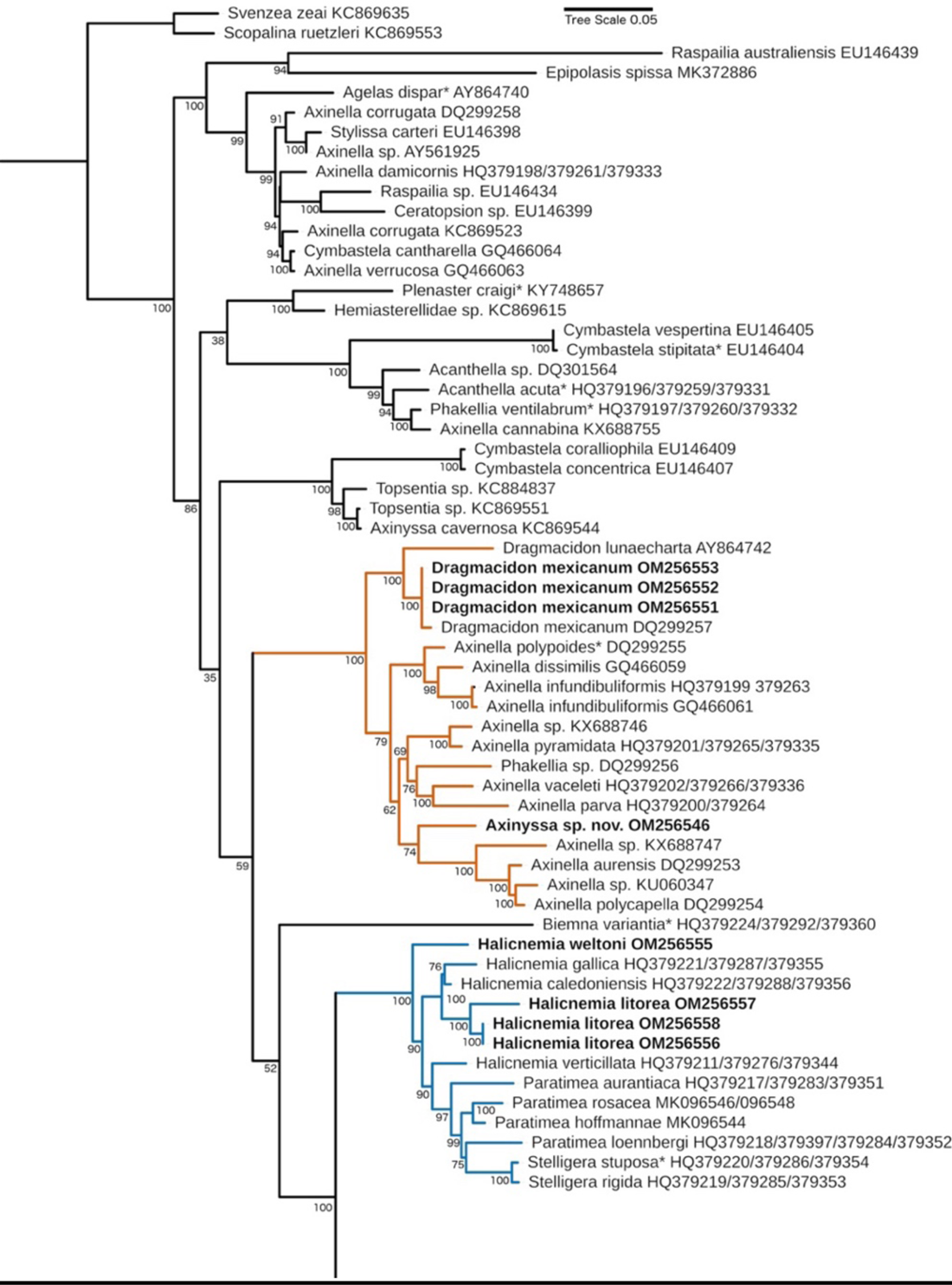
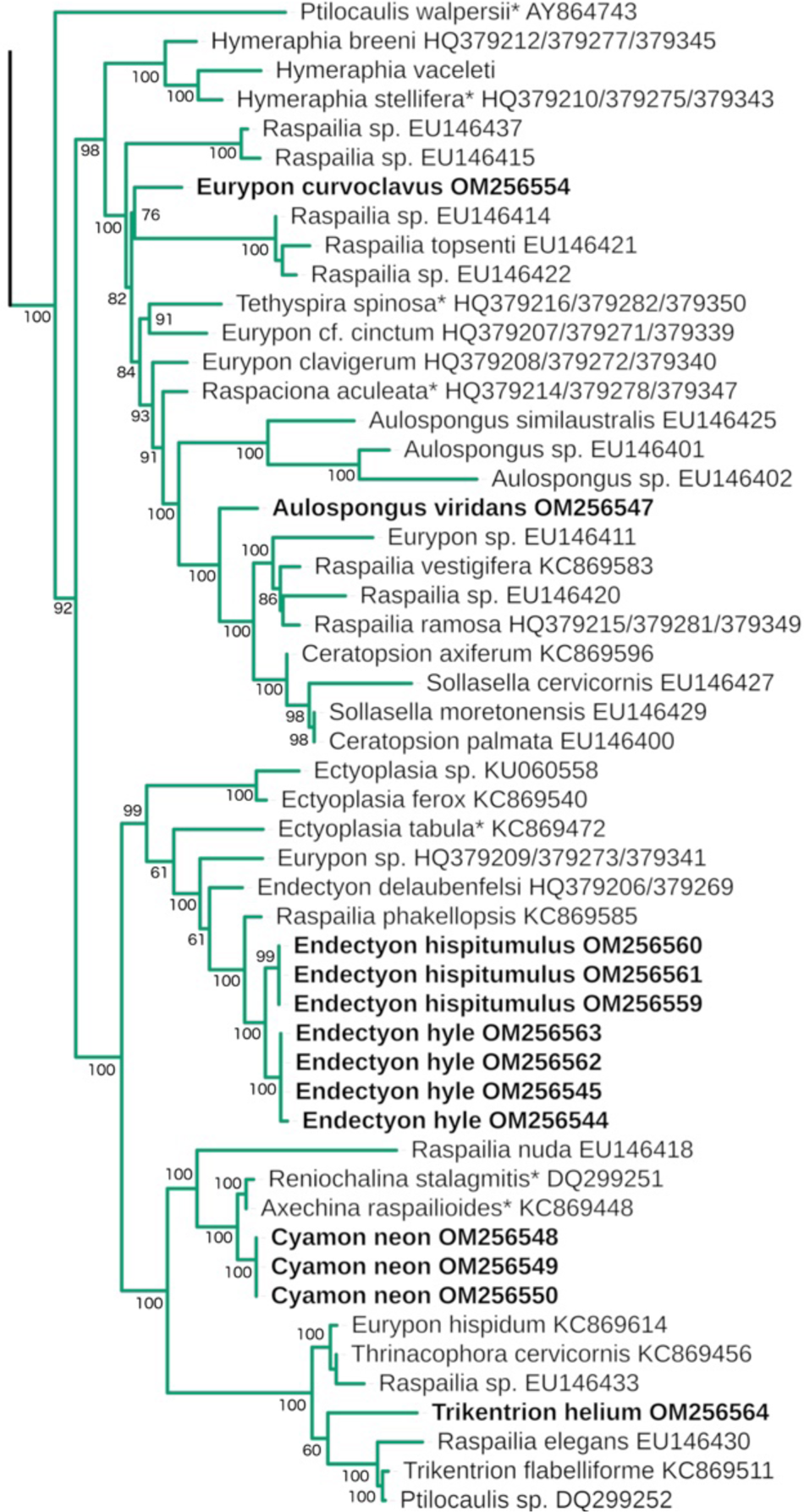
ML phylogeny of the large ribosomal subunit (28S). Genbank accession numbers are shown; bold indicates new sequences; asterisks designate type species. Node confidence is based on bootstrapping. Scale bar indicates substitutions per site. Root placement is based on Redmond et al. (2013). Orange = “Axinellidae *sensu stricto*”, Blue = Stelligeridae, Green = paraphyletic clade of mainly Raspailiidae.

The family Axinellidae contains 12 genera; our phylogeny includes species from seven of these, including the type species from five. Consistent with previous results, we find the family to be highly polyphyletic (Carballo *et al*. 2018; Gazave *et al*. 2010; Morrow *et al*. 2012; Redmond *et al*. 2013; Thacker *et al*. 2013). The type species of *Ptilocaulis* and *Reniochalina* fall within a clade, shown in green, that is mostly comprised of the Raspailiidae. The type species from *Phakellia* and *Cymbastela*, along with a subset of *Axinella* species, fall into several other well supported clades (shown in black) that include type species from other orders like the Agelasida and Bubarida.

The California species *Dragmacidon mexicanum,* however, falls within a clade, shown in orange, that includes the type species *Axinella polypoides* (Schmidt 1862). The presence of this species in this “Axinellidae *sensu stricto*” clade indicates that it will likely remain in this family after future revisions reconcile the phylogeny and taxonomy. Sequences from *D. mexicanum* form a clade with the only other *Dragmacidon* with data at this locus. In a separate phylogeny of the D3-D5 region of 28S (not shown), all 5 species of *Dragmacidon* with available data fell into this exclusive clade, so this genus may prove to be less polyphyletic than *Axinella*.

An additional member of this “Axinellidae *sensu stricto*” clade was found in California, shown in figure 1 as *Axinyssa* sp. nov. Morphological taxonomy places this species is in the genus *Axinyssa*, which is housed in the family Halichondriidae in the order Suberitida. This species was therefore not included in the current revision of the Axinellida, and will be described in a later revision of the Suberitida. We include this species in the phylogeny for completeness, as future revisions of the Linnean taxonomy are expected to result in this species moving to the Axinellidae. Existing data support the hypothesis that the genus *Axinyssa* is highly polyphyletic, with some species, including the type species, falling among the Bubarida and others among the Biemnida (Morrow *et al*. 2013; Morrow & Cárdenas 2015; Pankey *et al*. 2022; Thacker *et al*. 2013).

The family Stelligeridae (fig. 1, in blue) was recently revised by Morrow et al. (2019) using a combination of DNA and morphological data. As such, it now forms a monophyletic group with the exception of *Plenaster craigi*. Of the three new species of California stelligerids described below, we were able to obtain 28S sequence from two. Morphologically, both species are in the genus *Halicnemia*, which is polyphyletic based on this single locus. As data is only available from a few species thus far, not including the type species *H. patera* (Bowerbank 1864), revising this genus awaits further data.

Sequences are unexpectedly variable for *Halicnemia litorea* sp. nov., with pairwise differentiation of 4.6% between two isolates collected within a few kilometers of each other. The 28S locus is multicopy, and though gene conversion homogenizes these copies, there can sometimes be intragenomic polymorphism within species (Simon & Weiss 2008). This may be responsible for the apparently high variability among samples of this species, as one of the three individuals sequenced had a large number of bases that appeared to be variable in the raw sanger sequence data, and many of these variable sites were the same as the sites that differentiated the other two individuals.

The family Raspailiidae is large, with 5 subfamilies and 26 genera. New and published 28S sequences from the family largely fall within a single well-supported clade (shown in green), containing representatives of 13 of these genera. This paraphyletic clade also includes a few species currently placed in other families within the Axinellida (*Ptilocaulis*, *Reniochalina*), or in other orders (*Tethyspira* (Bubarida: Dictyonellidae); *Epipolasis* (Suberitida: Halichondriidae), *Ceratoporella* (Agelisida: Astroscleridae). Data from the mitochondrial cox1 locus tell a similar story, though the relationships among some Raspailliidae sub-clades are different; these data are shown in figure 2, with several Axinellidae *sensu stricto*, including new sequence from *D. mexicanum*, as outgroups. All six of the sequenced Raspailiidae from California fall within this large raspailiid clade at both loci, supporting their inclusion in the family.

**Figure 2.**
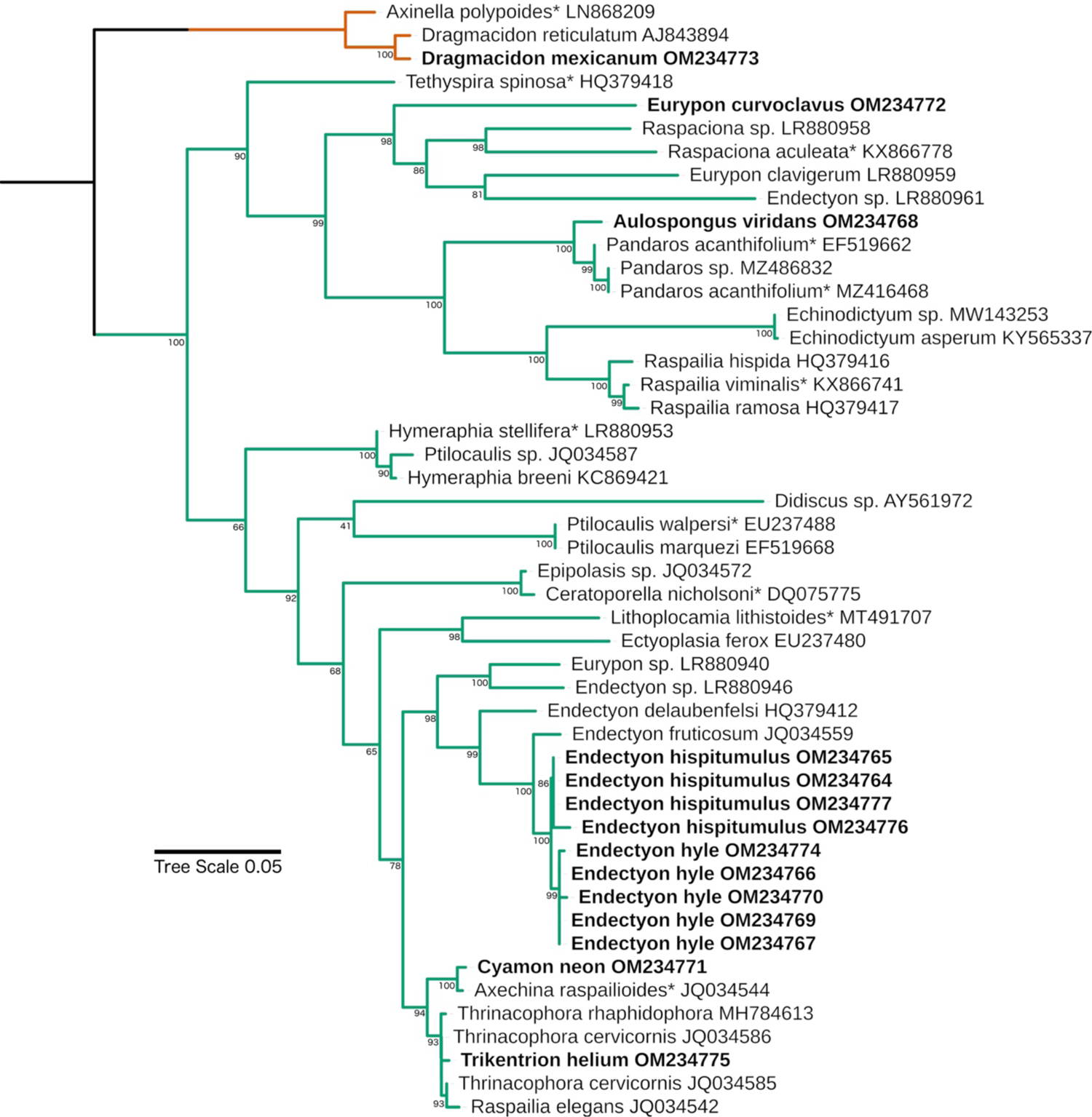
ML phylogeny of the cox1 mitochondrial locus. Genbank accession numbers are shown; bold indicates new sequences; asterisks designate type species. Node confidence is based on bootstrapping. Scale bar indicates substitutions per site. Root placement is based on Redmond et al. (2013). Green = paraphyletic clade of mainly Raspailiidae; Orange = subset of “Axinellidae *sensu stricto*” sequences used as outgroups.

**Figure 3.**
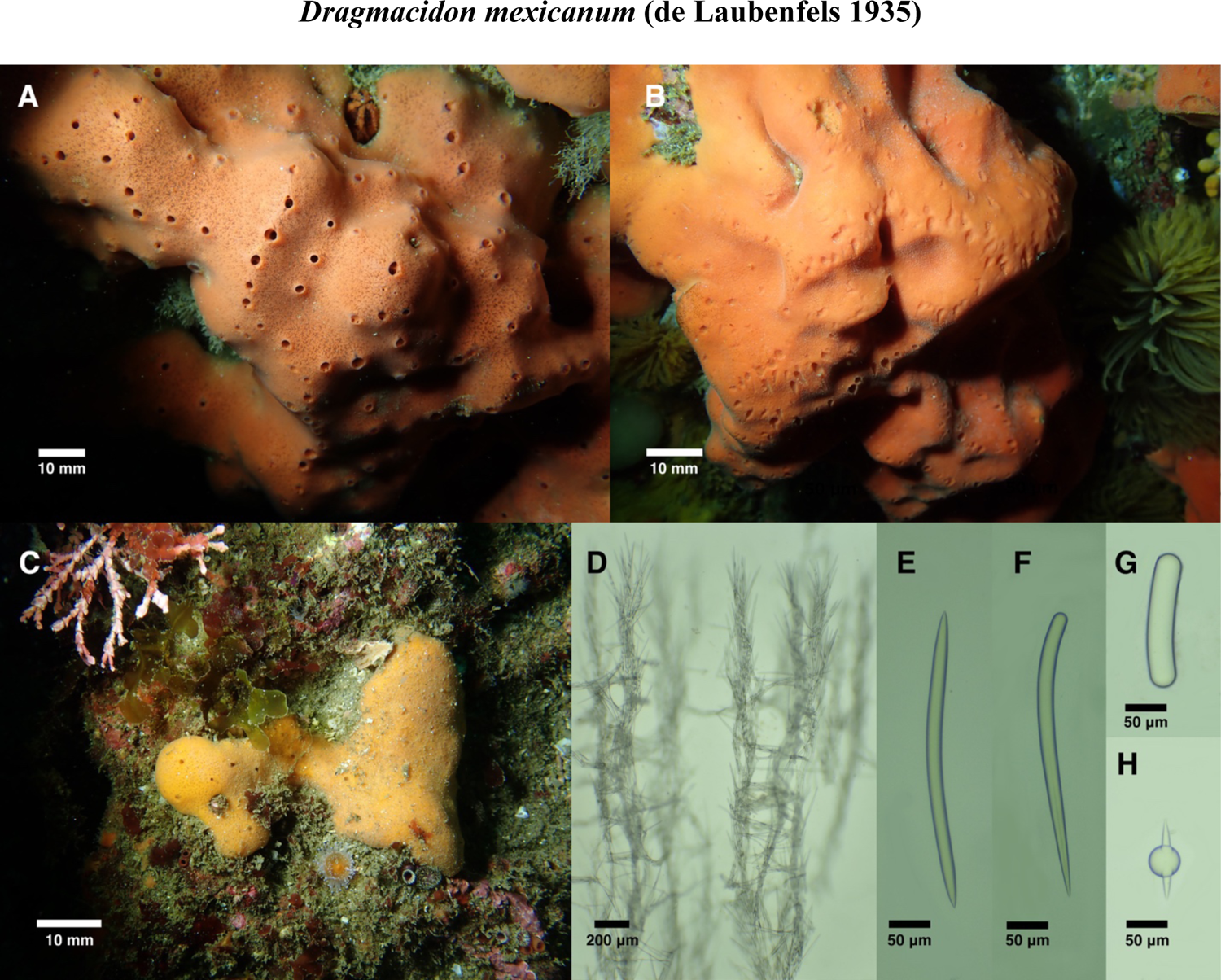
*Dragmacidon mexicanum*. A: Field photo of UCSB-IZC00041054, pores and oscules open. B: Field photo of SBMNH 693080, pores and oscules closed. C: Field photo of rare yellow individual SBMNH 693064, pores and oscules open for left portion only. D: Skeletal structure of UCSB-IZC00041054, brushes at top pierced surface of sponge. E-F: Oxea and style from SBMNH 693086. G: Capsule from SBMNH 693080. H: Sphere from SBMNH 693073.

In addition to the paraphyletic inclusions mentioned above, many raspailiid subfamilies, genera, and subgenera are polyphyletic. This is consistent with previous analyses (Erpenbeck *et al*. 2007b; Thacker *et al*. 2013), but our expanded dataset adds weight to this conclusion and offers additional insights. For example, the raspailiid subfamily Cyamoninae was previously represented only by *Trikentrion flabelliforme*. Surprisingly, this species clustered with members of the Thrinacophorinae at both 28S (Erpenbeck *et al*. 2007b) and 18S (Redmond *et al*. 2013). We have added two California Cyamoninae, *Cyamon neon* and *Trikentrion helium*: like *T. flabelliforme*, these species have polyactine megascleres not found in other subfamilies, and rare outside the order Tetractinellida (van Soest *et al*. 2012). Though the three species fall into the same clade, clustered with Thrinacophorinae and Raspailiinae, they do not form an exclusive clade with each other. This result is highly supported at both the 28S and cox1 loci and therefore appears to be a legitimate reflection of the species tree. It would be surprising if polyactine megascleres had evolved multiple times in the Raspailiidae; the alternative explanation of multiple losses may be more likely.

It is also notable that the newly generated cox1 sequence from *Trikentrion helium* (from California), is identical to sequences of both *Thrinacophora rhaphidophera* (from Vietnam) and *Thrinacophora cervicornis* (from Australia) across the entire Folmer barcoding region (∼650 bp; 1150 bp of cox1 were sequenced for *T. helium*, but only the Folmer region is available from the other species). This cannot be due to contamination, as no Raspailiidae, other than those sequenced in the current paper, have ever been present in the lab where the current work was done. As the morphology of these species is quite distinct, and the sequences at the 28S locus well differentiated, this adds to a list of examples of a slow rate of evolution at the cox1 locus in sponges (Huang *et al*. 2008; López-Legentil *et al*. 2010; Pöppe *et al*. 2010).

Five species of *Eurypon* are represented in the 28S tree, including *E. curvoclavus* sp. nov. All five species are separated from each other in the phylogeny, emphasizing that the genus, as currently defined, is not tenable. As there are over 50 species in the genus, acquiring genetic data from more species — especially the type species, *E. clavatum* — would be helpful in forming new hypotheses regarding the evolution of encrusting raspailiids.

The newly sequenced *Aulospongus viridans* sp. nov. does not form a clade with other *Aulospongus* at 28S, but with only a few members sequenced, and no data from the type species, firm conclusions regarding this genus seem premature. Interestingly, at cox1, *A. viridans* is closely related to the monotypic *Pandaros acanthifolium*. This species is currently placed in the Microcionidae (order Poecilosclerida), but, as noted in the Systema Porifera, “*Pandaros* is a borderline taxon that could be legitimately included in either Raspailiidae or Microcionidae” (Hooper 2002b). Though *A. viridans* shares lightly spined, slightly rhabdostyle-shaped styles with *P. acanthifolium*, the new species does not fit the other important features of the genus *Pandaros* as defined by Hooper (2002). It seems possible that *A. viridans* will be included in a revised version of the genus *Pandaros* in the future, once more species have been sequenced and the defining features of this clade have been determined. In the meantime, we suggest that *Pandaros* be moved to the family Raspailiidae.

Two newly sequenced *Endectyon* are the only members from the California Axinellida that appear closely related to one another. The genetic data provide strong evidence that they are indeed sister species, and not morphotypes of the same species, because there are fixed genetic differences at both 28S and cox1. Because nuclear and mitochondrial genes are unlinked, even low amounts of gene flow between them should disrupt such a pattern; because these species are sympatric, this indicates that other types of reproductive isolation separate them.

These California *Endectyon* also provide an interesting perspective on the phylogenetic signal of acanthostyle spine morphology. The two California *Endectyon* form a clade with *Endectyon (Endectyon) delaubenfelsi* at 28S, and both *E. (E.) delaubenfelsi* and *E. (E.) fruticosum* at cox1. Also included in this clade is *Raspailia (Raspailia) phakellopsis*: the description of this species shows large, recurved hooks on the acanthostyles (Hooper 1991). This is a trait shared by *Endectyon* but absent from the other named species in the phylogeny. The only exception is *Ectyoplasia*, which have similar recurved spines, and at 28S, this clade is sister to the *Endectyon* clade. These results may indicate that the type of spination on acanthostyles is more phylogenetically informative than previously realized. In contrast, the loss of acanthostyles has most likely happened repeatedly. Species without acanthostyles often grouped with other species without acanthostyles, but these groups are dispersed throughout the phylogeny (e.g. [*Sollasella* + *Ceratopsion*], [*Axechina* + *Reniochalina*]).

### Systematics

#### Summary of California Axinellida

Family Axinellidae (Carter 1875)
  *Dragmacidon mexicanum* (de Laubenfels 1935)

Family Raspailiidae (Nardo 1833)
  Subfamily Raspailiinae (Nardo 1833)

*Endectyon (Endectyon) hyle* comb. nov. (de Laubenfels 1930)
*Endectyon (Endectyon) hispitumulus* sp. nov.
*Raspailia (Raspaxilla) hymani* (Dickinson 1945)
*Eurypon curvoclavus* sp. nov.
*Aulospongus viridans* sp. nov.
*Aulospongus lajollaensis* sp. nov.

Subfamily Cyamoninae (Hooper 2002c)
  *Cyamon neon* (de Laubenfels 1930)
  (*Cyamon argon* (Dickinson 1945) placed in synonymy with *C. neon*)
  *Cyamon koltuni* (Sim & Bakus 1986)
  *Trikentrion helium* (Dickinson 1945)
  (*Trikentrion catalina* (Sim & Bakus 1986) placed in synonymy with *T. helium*)
  Family Stelligeridae (von Lendenfeld 1898)
    *Halicnemia litorea* sp. nov.
    *Halicnemia montereyensis* sp. nov.
    *Halicnemia weltoni* sp. nov.

**Family Axinellidae (Carter 1875)**

**Genus Dragmacidon (Hallmann 1917)**

Axinella mexicana (de Laubenfels 1935)

Axinella sp. (Preston *et al*. 1996)

Pseudaxinella mexicana (Gómez *et al*. 2002)

Dragmacidon mexicanum (Sim & Bakus 1986)

?Dragmacidon sp. (Lee *et al*. 2007a)

### Material examined

UCSB-IZC00041064 and TLT16, Elwood Reef, Santa Barbara (34.41775, - 119.90150), 9-15 m, 4/17/2019; SBMNH 693101, Otter Cove, Monterey (36.62920, −121.92031), 7-12 m, 11/24/2019; UCSB-IZC00041054, UCSB-IZC00041055, UCSB-IZC00041057, SBMNH 693057 & SBMNH 693058, Big Rock, Santa Cruz Island (34.05220, − 119.57360), 9-14 m, 1/19/2020; SBMNH 693073 and UCSB-IZC00041059, Goalpost, Point Loma, San Diego (32.69438, −117.26860), 12-15 m, 2/8/2020; SBMNH 693064, Dino Head, Point Loma, San Diego (32.68808, −117.27080), 21-26 m, 9/19/2020; TLT97, Mohawk Reef, Santa Barbara (34.39407, −119.72957), 5-11 m, 6/28/2019; SBMNH 693072, Farnsworth Bank, Catalina Island (33.34380, −118.51650), 21-27 m, 7/10/2021; SBMNH 693063, CRABS, Monterey (36.55377, −121.9384), 10-17 m, 9/21/2021; SBMNH 693081 and SBMNH 693080, Honeymooners, Carmel Bay, Monterey (36.50390, −121.94100), 10-20 m, 9/22/2021; SBMNH 693061, Cannery Row, Monterey (36.61798, −121.8978), 9-16 m, 9/21/2021; SBMNH 693085, TLT142, TLT238 & TLT241, Naples Reef, Santa Barbara, California (34.42212, −119.95154), depth 10-15 m, 7/31/2019; TLT63, TLT64, TLT111 & TLT112, Landslide Cove, Anacapa Island (34.01539, −119.43560), 7-11 m, 4/25/2019, collected by SP & TT; TLT53 & TLT73, West End, Anacapa Island (34.01352, −119.44570), 7-11 m, 4/25/2019, collected by SP & TT; SBMNH 693086, Naples Reef, Santa Barbara (34.42212, −119.95154), depth not recorded, 7/15/2019, collected by Robert Miller; UCSB-IZC00041050 & UCSB-IZC00041051, Prince Island (near San Miguel Island), California (34.056995, −120.331089), depth not recorded, 10/8/2019, collected by Frankie Puerzer and Robert Miller; NHMLA 15927 / BULA-0238, Marineland, Palos Verdes, Los Angeles (33.75166667, −118.41665), depth not recorded, collected by Gustav Paulay. Samples collected by TT except where noted.

### Morphology

Thickly encrusting to massive, 5 mm to 5 cm thick. One of the largest shallow-water sponges in the region, with some over 60 cm across. Most are smooth, thick encrustations, 10-20 mm thick. Some are flat and even, others are irregular in thickness, with low undulating mounds and variable intensity of pigment. The thickest specimens are up to 5 cm thick, massive and irregular, with prominent ridges. Nearly all are orange or orange-red externally and internally for the first 1-3 mm, and yellow internally; a few are yellow externally and internally. All sponges fade to beige in ethanol. Surface with numerous oscules 1-2 mm in diameter, flush with surface or slightly elevated; in massive sponges, oscules occur primarily along ridges and other high points. Sponge surface between oscules is prominently patterned with a mix of opaque and transparent tissue; dermal pores (each 50-100 μm diameter) visible in transparent regions. Oscules are closeable with a membrane, and sponges are frequently found with oscules and pores closed, in which case they appear entirely opaque and without surface patterning, often with sediment accumulating on the surface. Surface appears smooth to the eye but is microscopically hispid. Produces copious slime when cut.

#### Skeleton

Plumose ascending columns of styles and oxeas, chaotically bridged by oxeas. Columns pierce surface of sponge to create microscopically hispid surface.

#### Spicules

Styles and oxeas.

Styles: usually curved or bent near head end. Size is variable among individuals, with mean length varying from 263 to 371 μm (n=15 sponges, n=8-27 spicules per sponge); pooled data from all sponges 186-318-441 x 4-13-23 μm (n=315).

Oxeas: usually bent in the center. Mean length per sponge varies from 315 to 380 μm (n=15 sponges, n=7-26 spicules per sponge); pooled data from all sponges 255-351-439 x 3-13-23 μm (n=272). Oxeas are longer than styles in 14 of 15 samples, with oxea/style length ratio varying from 0.96-1.16.

Spheroids: one sample (SBMNH 693073) had spherical and capsule-shaped spicules, 21-65 μm in diameter, some with one or two protruding spikes. These are interpreted as an aberration.

### Distribution and habitat

Common on subtidal rocky reefs throughout the investigated region (Central and Southern California) and previously known from Baja California and the Sea of Cortez (Gómez *et al*. 2002). This species was found at all subtidal depths investigated (5-30 m), but has not been found in the intertidal zone, or on any human-made structures (shipwrecks, oil rigs, pier pilings, or floating docks). It is especially abundant in kelp forests around Point Loma and La Jolla in San Diego County, where it was seen at all 8 sites searched. It was seen at 18 of 21 natural reefs at Anacapa and Santa Cruz Islands, but only 4 of 13 natural reefs on the mainland side of the Santa Barbara Channel. This could be due to a preference for warmer water, but based on where it was found on the mainland, we suspect it also indicates a preference for sites with lower suspended sediment. It was also common in the colder waters of Central California, as it was found at 6 of 14 subtidal sites around the Monterey Peninsula.

## Discussion

This common, large, and brightly colored species is one of the most apparent subtidal sponges in Southern and Central California, and its archaeal symbionts have been a focus of past work (Preston *et al*. 1996). Nonetheless, the identity of this species was previously uncertain. When reviewing the literature on California sponges, Lee et al. (2007a) reported this species as “*?Dragmacidon sp.”* because the quantitative spicule dimensions of California sponges were different from samples described from Baja California (Dickinson 1945; de Laubenfels 1935). A more recent and detailed report of samples from the Sea of Cortez matches California sponges qualitatively and quantitatively (Gómez *et al*. 2002). Mexican samples are reported as having oxeas only 5% longer, and styles 1% shorter than the pooled means reported here. Mexican sponges are reported as having 19%-26% thicker spicules than California. The data reported here are consistent with considerable variability in spicule dimensions within populations, so it seems likely that the California population fits comfortably with the named species *D. mexicanum*. Though more genotyping could reveal cryptic species, we found no evidence of any within California. Genotyped samples include the sample with the shortest spicules and two with spicules of average size, and there was no association of spicule size with geographic location. Two other species of *Dragmacidon* are known from the Mexican Pacific and could range into California: *Dragmacidon oxeon* (Dickinson 1945) and *D. ophisclera* (de Laubenfels 1935). Both have much longer spicules and we know of no records matching their description from California. Likewise, *D. kishinense* (Austin *et al*. 2013) is known from British Columbia and could range into California, but has much longer spicules and no California records are known.

*Dragmacidon mexicanum* can be tentatively identified in the field based on gross morphology alone, but spicule or DNA confirmation is advisable due to the morphological similarity of several sympatric sponges. Chief among these is *Acarnus erithacus* (de Laubenfels 1927). These species can often be distinguished using the more prominent surface patterning of *D. mexicanum*, which appears as a mesh of small pores with a maze-like pattern of opaque orange flesh below; *A. erithacus* is entirely opaque or with faintly visible pores. The oscula also differ, on average, with *A. erithacus* displaying oscules atop large cone or chimney-shaped projections, while *D. mexicanum* has oscules flush with the surface or slightly elevated; in large *D. mexicanum* oscula occur along ridges but generally not atop individual cone-shaped projections. Also, *D. mexicanum* is firm but yielding, while *A. erithacus* is rock-hard. Some sympatric *Antho sp.* can obtain large size and then closely resemble *D. mexicanum*, especially around the Monterey Peninsula in Central California. Like *A. erithacus*, these *Antho sp.* generally lack the prominent surface patterning between oscula.

Clint Nelson, the Dive and Boat Safety Officer for the Santa Barbara Coastal Long-Term Ecological Research Site, reports using a specific, large *D. mexicanum* as an underwater navigational landmark for the past 16 years, so this species is likely to be long lived.

**Family Raspailiidae (Nardo 1833)**

**Subfamily Raspailiinae (Nardo 1833)**

**Genus Endectyon (Topsent 1920)**

Hemectyon hyle (Bakus & Green 1987; Dickinson 1945; Green & Bakus 1994; de Laubenfels 1932)

Aulospongus hyle (Desqeyroux-Faundez & van Soest 1997)

Raspailia (Raspaxilla) hyle (Aguilar-Camacho & Carballo 2013; Hooper *et al*. 1999) Endectyon (Endectyon) hyle (Lee *et al*. 2007a)

### Material examined

NHMLA 16545 / BULA-0541, Hawthorne Reef, Los Angeles, California, (33.74714, −118.42090), 24 m depth, 8/23/2019, collected by TT; SBMNH 693066, SBMNH 693067, SBMNH 693068 & UCSB-IZC00041048, Elwood Reef, Santa Barbara, California, (34.41775, −119.90150), 10-15 m depth, 10/23/2019, collected by TT; UCSB-IZC00041049, Prince Island (near San Miguel Island), California (34.056995, −120.331089), depth not recorded, 10/8/2019, collected by Frankie Puerzer and Robert Miller; UCSB-IZC00041044, Honeymooners, Monterey, California, (36.50390, −121.94100), 12-19 m depth, 9/22/21, collected by TT; UCSB-IZC00041042, Cannery Row, Monterey, California, (36.61798, − 121.8978), 9-15 m depth, 9/21/21, collected by TT; UCSB-IZC00041043, Inner Pinnacle, Monterey, California, (36.55852, −121.96820), 18-24 m depth, 9/21/21, collected by TT.

### Morphology

Growth upright but bushy, up to 6 cm in height. Bulky fronds extend upwards and outwards from a stalk or small base; each frond spreads along multiple axes and is usually covered in folds and corrugations. Some samples less frond-like and more lobe-like and columnar (figure 4B). Samples usually brownish-red when alive, but one sample was yellow; samples fade to tan in ethanol. Very hispid.

**Figure 4.**
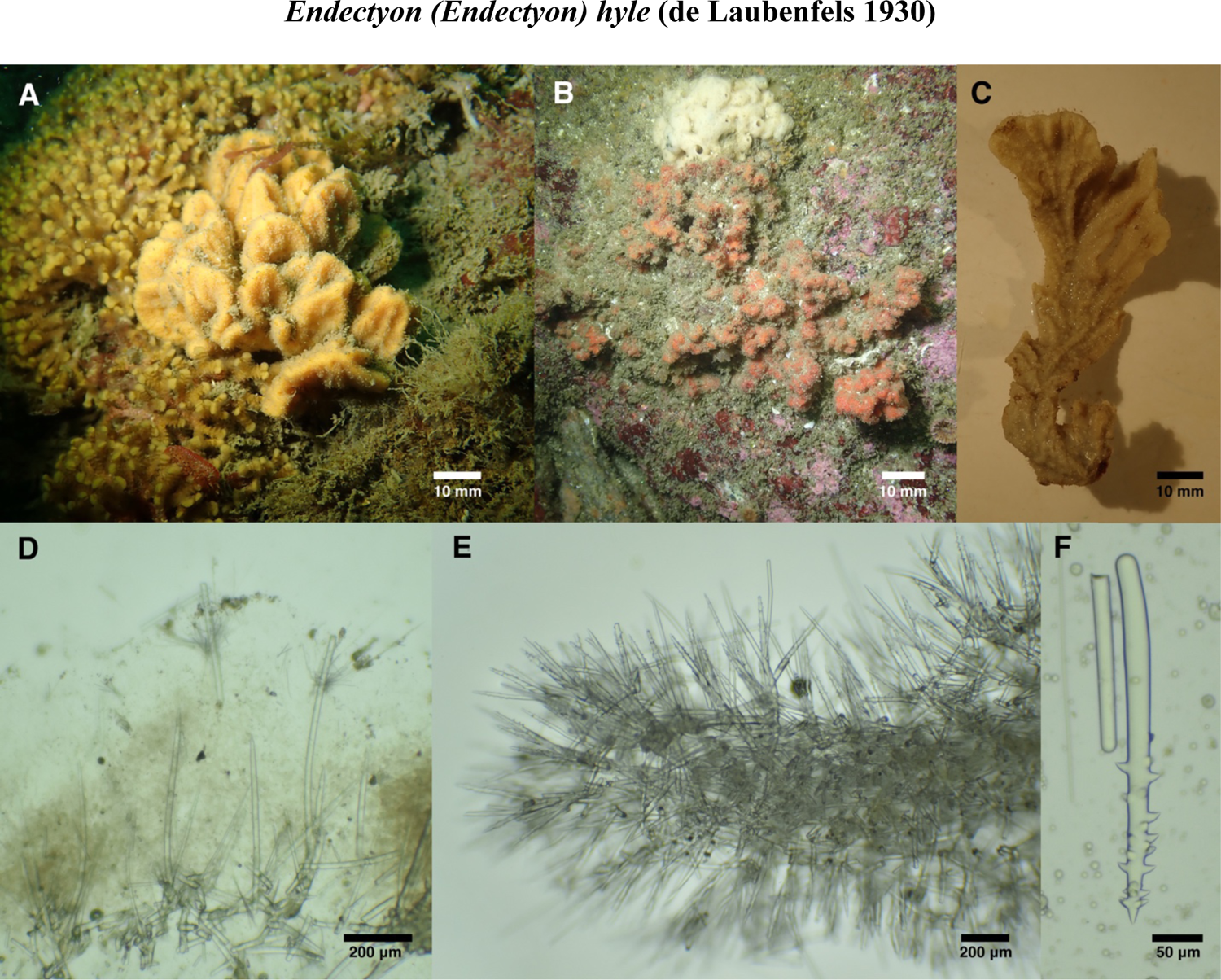
*Endectyon hyle.* Field photos of A: SBMNH 693068 and B: UCSB-IZC00041042. C: NHMLA 16545 post-preservation. D: Peripheral skeleton of UCSB-IZC00041048 showing long protruding styles and ectosomal bouquets of thin styles. E: Axial skeleton with echinating acanthostyles (NHMLA 16545). F: Typical acanthostyle (UCSB-IZC00041042).

**Figure 5.**
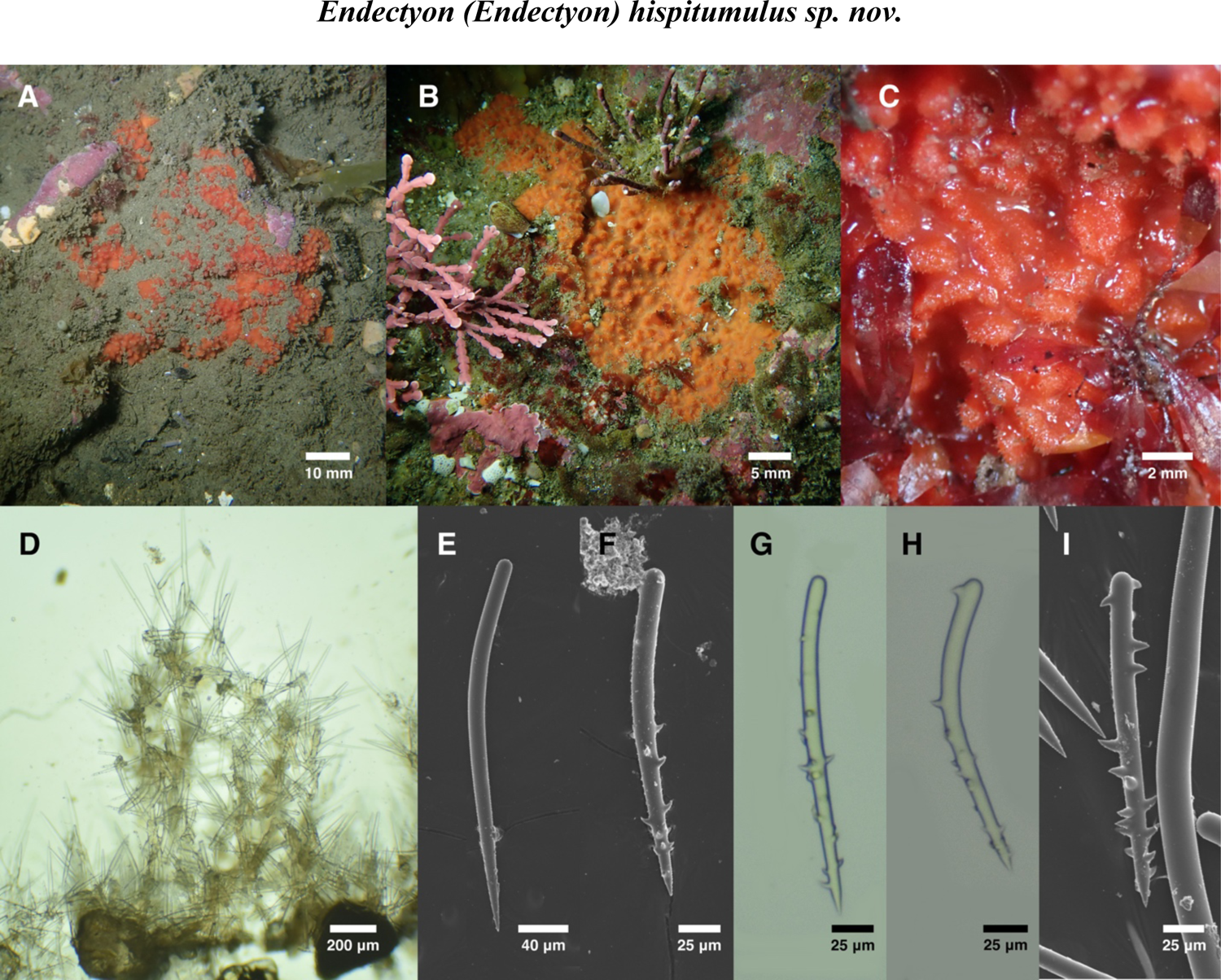
*Endectyon hispitumulus.* Field photos of A: holotype (CASIZ 236512), B: UCSB-IZC00041056; C: intertidal sample SBMNH 693084. D: Skeletal structure of UCSB-IZC00041056. E: Typical style from holotype. F-I: Spectrum of acanthostyle morphology; F, H from holotype, G from SBMNH 693074, I from UCSB-IZC00041056.

**Figure 6.**
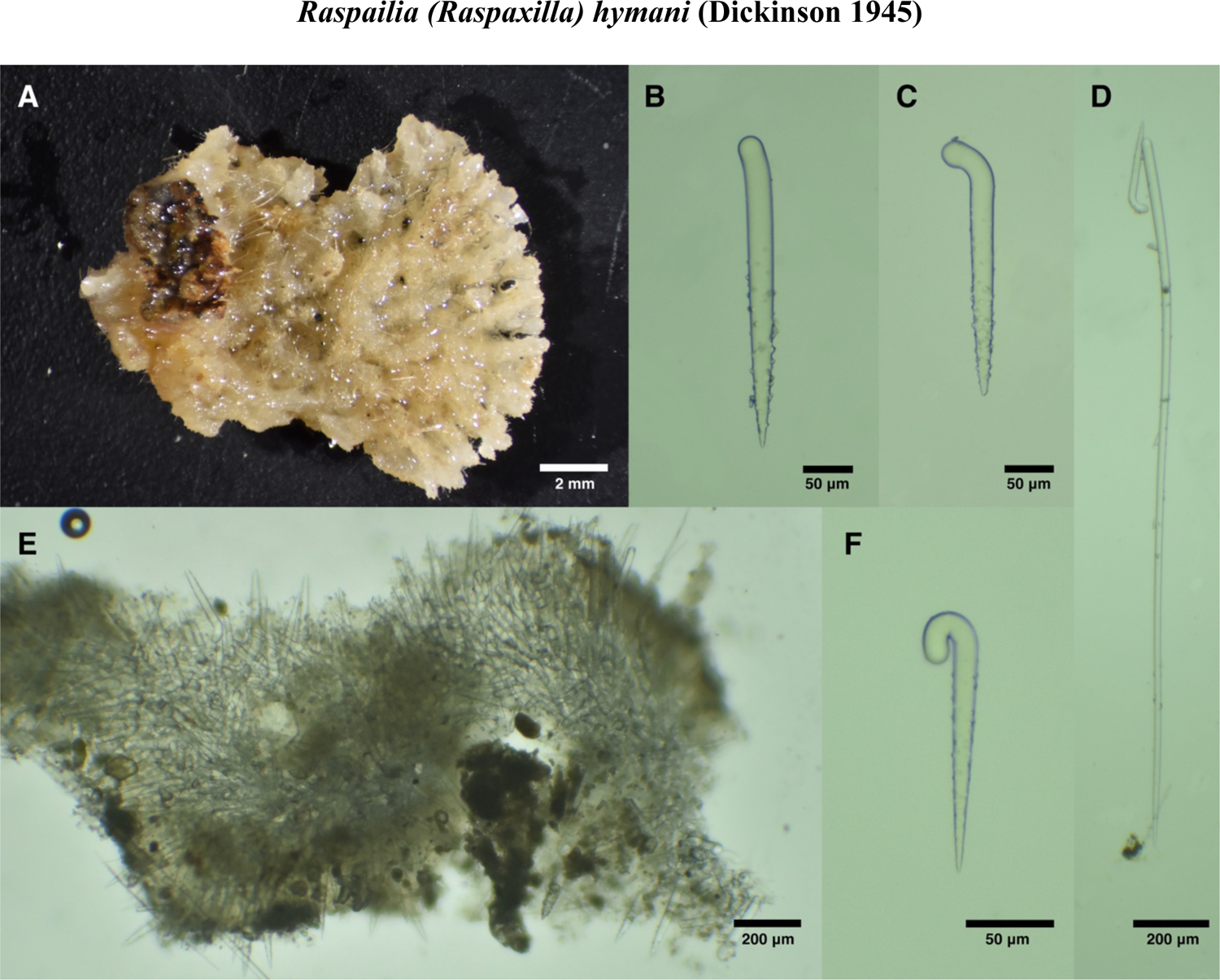
*Raspailia hymani.* A: Preserved sample. B, C, F: Spectrum of acanthostyle morphology. E: Skeletal structure. D: Long style with acanthostyle. All photos of sample SBMNH 467982.

**Figure 7.**
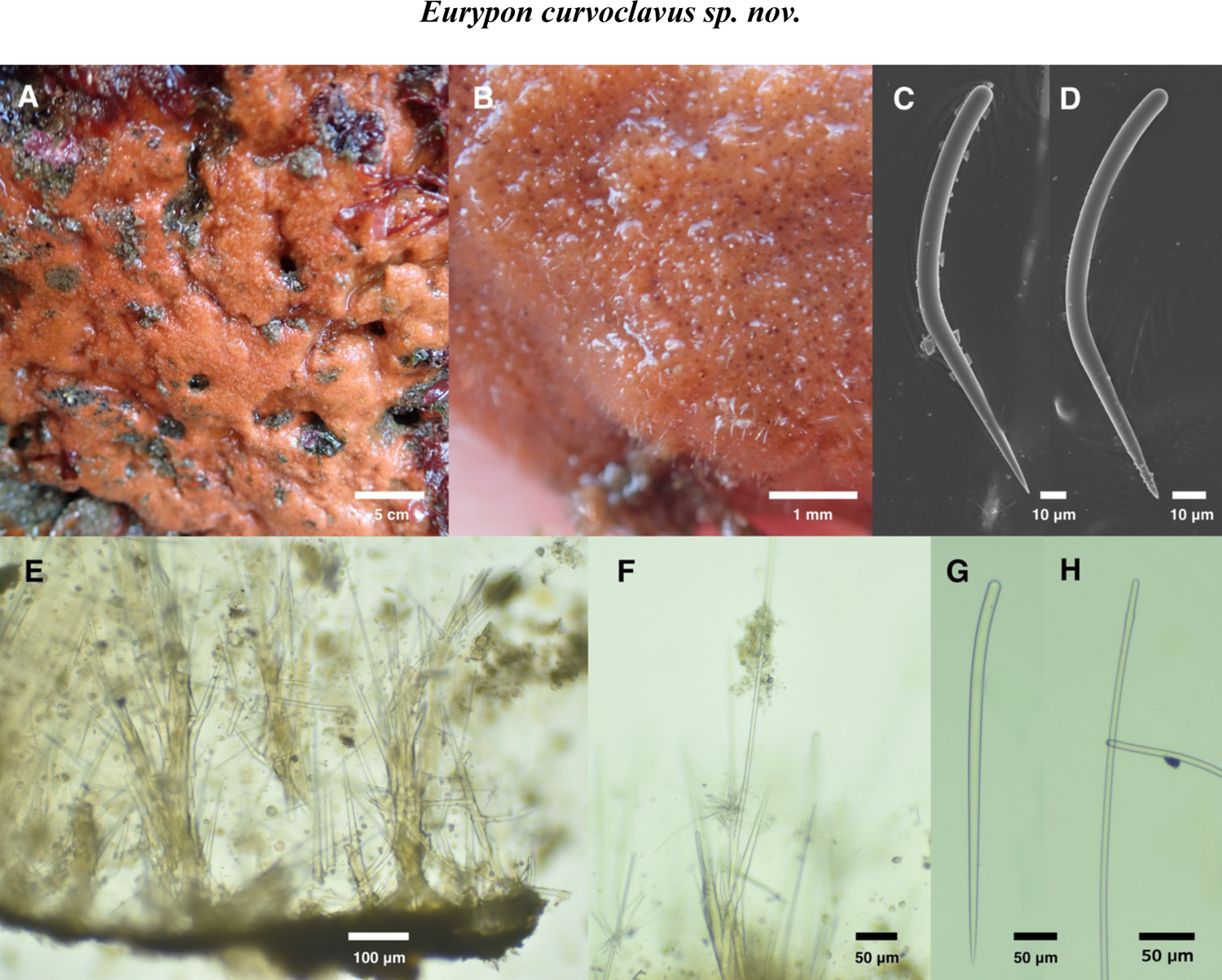
Field photos of A: holotype (CASIZ 236513) and B: UCSB-IZC00041061, both exposed at low tide. C: Acanthostyle without spines. D: Acanthostyle. E: Cross section showing spongin nodes arising from basal plate. F: Peripheral skeleton with long protruding style surrounded by ectosomal bouquet. G: Style. H: Long style. C-H from holotype.

**Figure 8.**
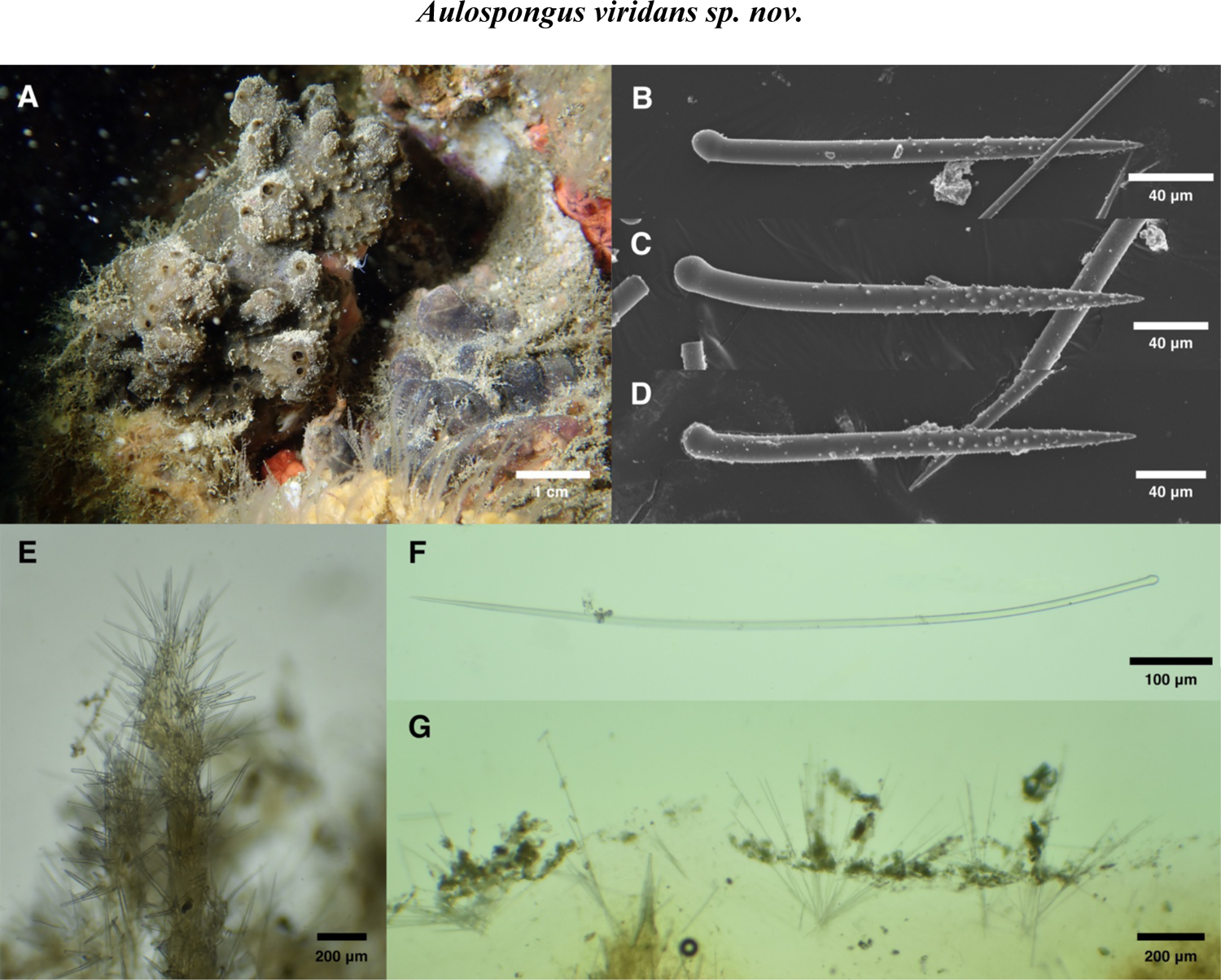
*Aulospongus viridans*. A: Field photo. B: Spectrum of morphology found in acanthostyles. C: Skeletal structure. D: Long subectosomal style. E: Ectosomal bouquets and surface debris. All photos from holotype, CASIZ 236515 / UCSB-IZC00041063.

**Figure 9.**
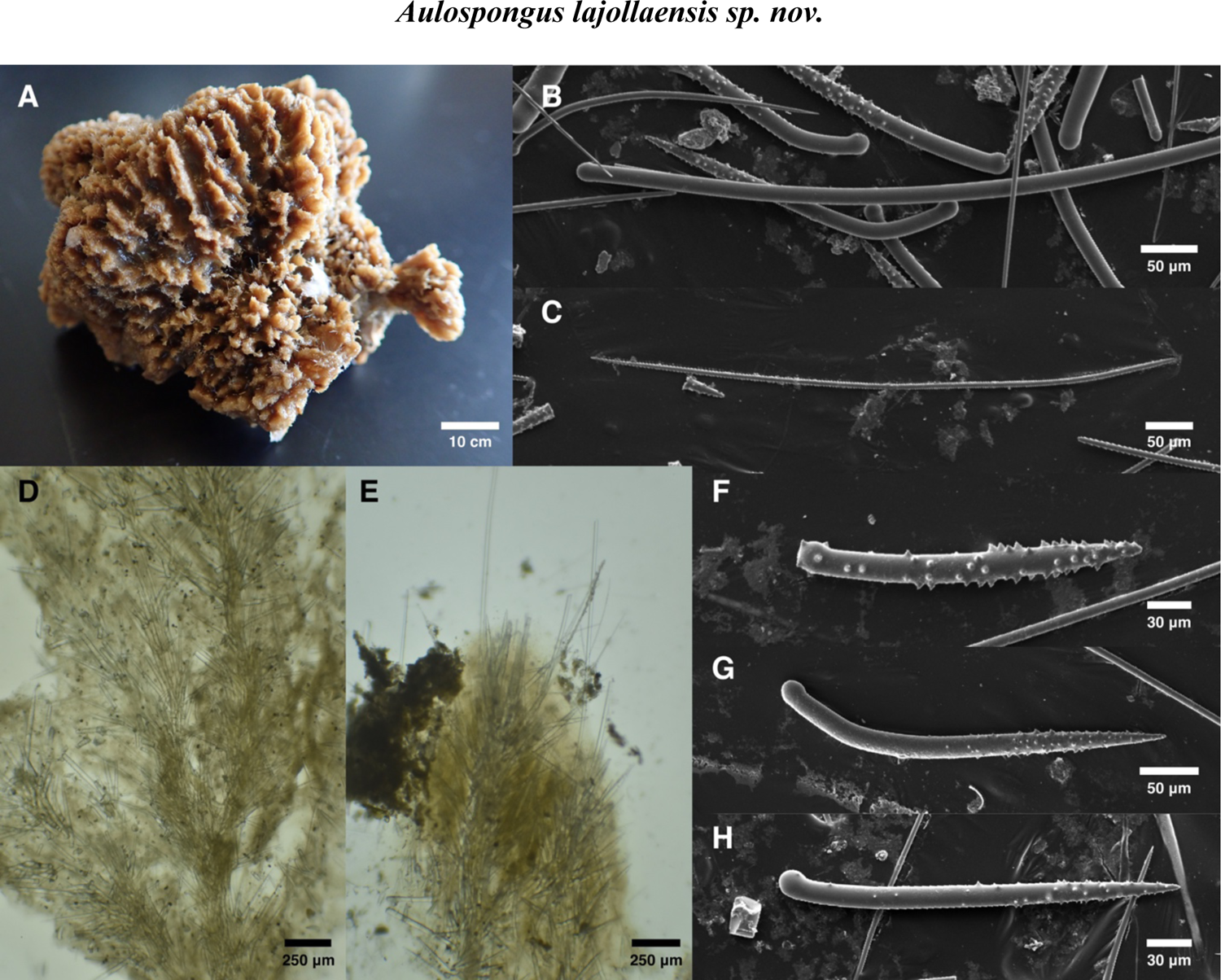
*Aulospongus lajollaensis*. A: Holotype post-preservation. B: Head of long subectosomal style in center, with rhabdostyles behind; thin, unspined rhabdostyle also visible in top left. C: Ectosomal oxea. D: Choanosomal skeleton. E: Cross-section showing protruding subectosomal styles and ectosomal oxeas. F: Atypical rhabdostyle. G-H: Typical rhabdostyles.

**Figure 10.**
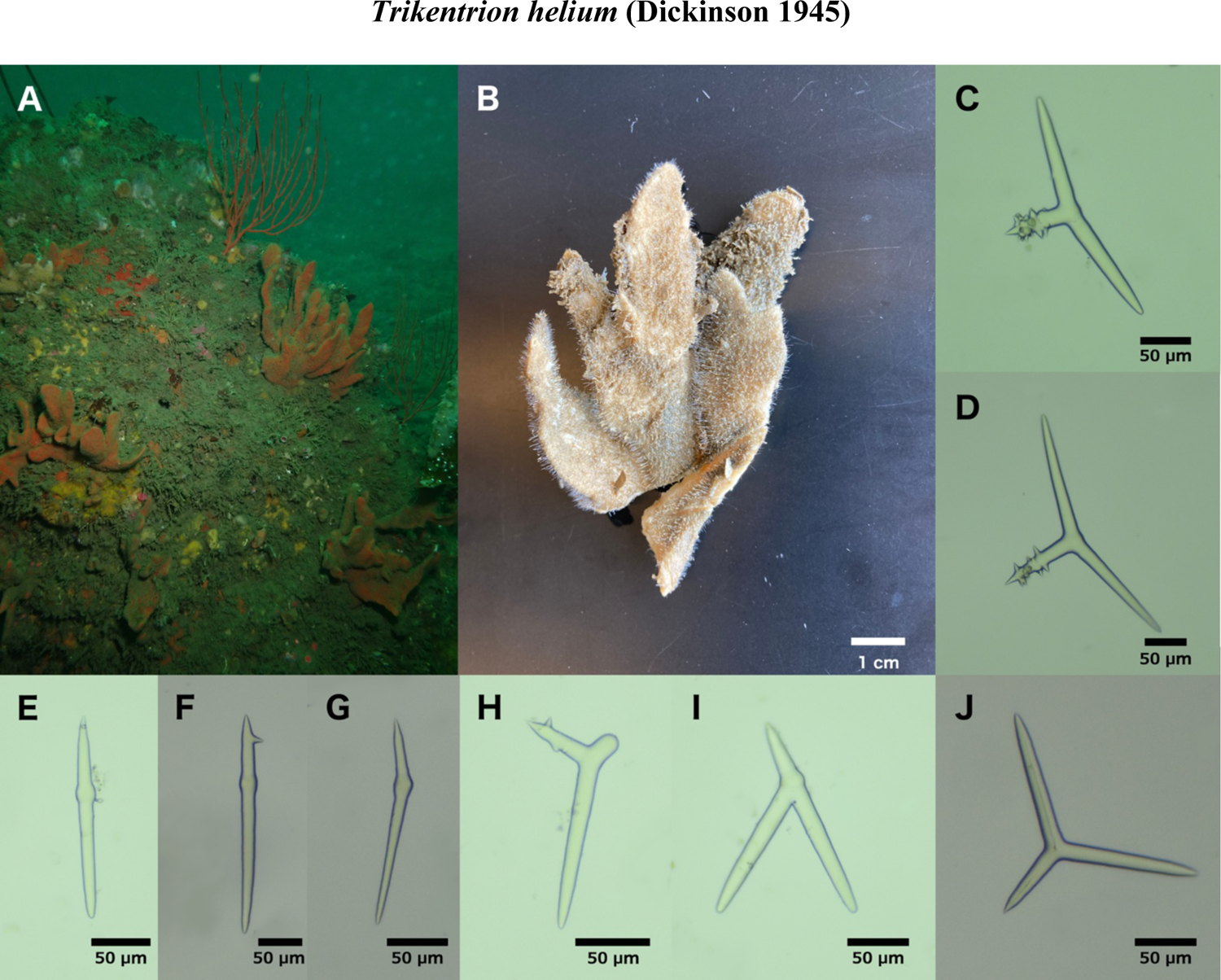
*Trikentrion helium.* A: Field photo of uncollected samples off Point Loma, San Diego, 25 m depth. B: Sample NHMLA 20362 post-preservation. Typical triactines from UCSB-IZC00041058 (C) and NHMLA 20362 (D). Additional polyactine morphologies from UCSB-IZC00041058 (I) and NHMLA 20362 (H, J). Sampling of diactinal polyactines from UCSB-IZC00041058 (E) and NHMLA 20362 (F, G).

**Figure 11.**
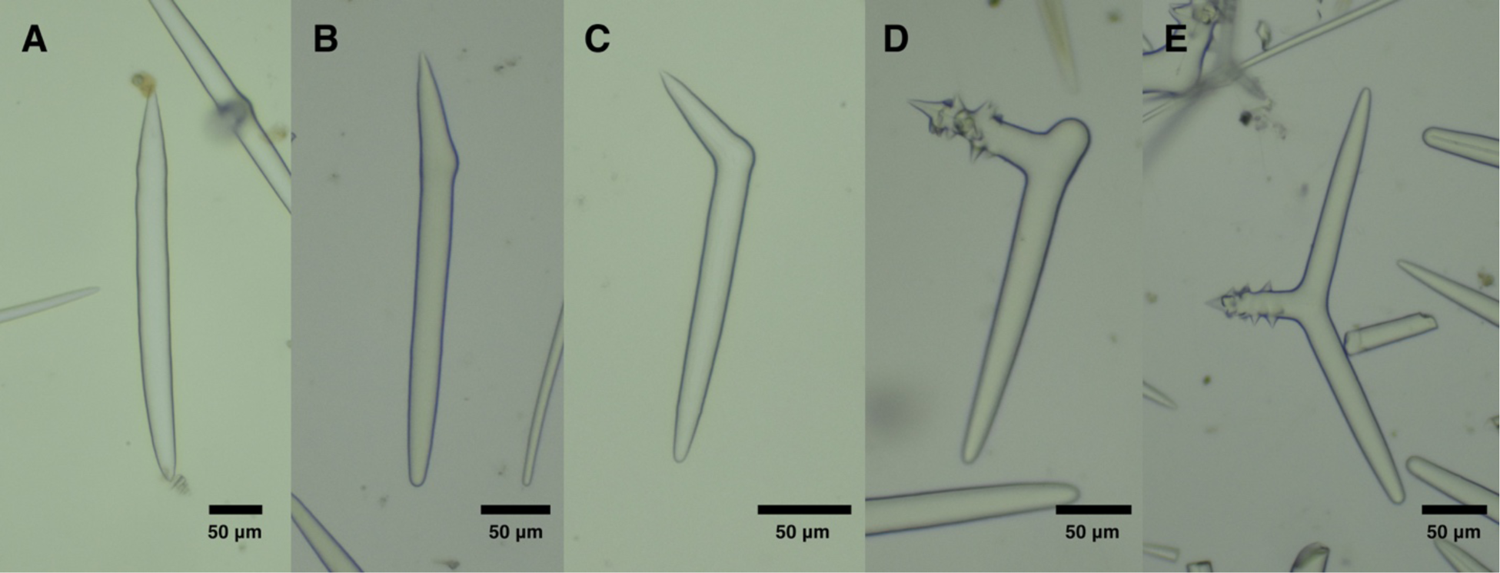
Polyactine morphologies from USNM 3361, the *Trikentrion catalina* holotype. A-C: diactinal polyactines. D: polyactine with one clade reduced to knob. E: typical triactine.

### Skeleton

Axially-compressed region is a dense reticulation of styles; some acanthostyles are within this reticulation, but most are echinating (surrounding reticulation, with heads imbedded in spongin and tips pointing outward). Long styles surround the axial reticulation much like acanthostyles, but unlike acanthostyles, they pierce the sponge surface to create hispidity. Very thin styles form ectosomal bouquets in typical raspailiid fashion.

### Spicules

Short styles, long styles, acanthostyles, and thin ectosomal styles; oxeas/strongyloxeas/strongyles sometimes present. Short styles, long styles, and acanthostyles are each 13-22% longer in Central California samples than Southern California samples.

Short styles: unadorned or with weak terminal or subterminal swelling. Often curved or bent, with the bend most commonly near the head end but often in the middle; Southern California: 281-454-582 x 3-12-24 μm (n=83), Central California: 385-520-682 x 8-18-26 μm (n=102).

Long styles: usually unadorned but sometimes with weak terminal or subterminal swelling; straight or curved; Southern California: 768-1065-1411 x 9-12-16 μm (n=10), Central California: 1016-1370-1791 x 12-20-27 μm (n=24).

Acanthostyles: occasionally bent near head but more often straight or slightly curved; spines usually limited to the half of style closer to the point, but sometimes occur near or even on the head of the style. Spines large and curved towards head end of style; Southern California: 90- 278-394 x 2-14-20 μm (n=64), Central California: 199-343-471 x 12-21-30 μm (n=62).

Thin ectosomal styles: Southern California: 226-350-768 x 1-3-5 μm (n=20), Central California: 1162-376-611 x 2-3-9 μm (n=57).

Oxeas/strongyloxeas/strongyles: present in some samples; dimensions similar to styles but often show two bends instead of one bend or a curve; Southern California: 502-572-657 x 8-13-17 μm (n=12), Central California: 585-753-868 x 13-21-26 μm (n=3).

### Distribution and habitat

Published occurrences range from British Columbia (Austin 1985) to the Gulf of California (Dickinson 1945), and from 15 m to 330 m in depth (Luke 1998). We found this species only rarely on shallow (<15 m) rocky reefs, where most search effort was concentrated, but more frequently on deeper (24-30 m) reefs. It was found in multiple previous surveys around Southern California, with some samples reportedly as shallow as 15 m (sample CASIZ 182375, Santa Rosa Island) but most “shallow” samples were collected below 20 m (Luke 1998). It has been found in multiple deep-water surveys far below diving depths in Southern California (Bakus & Green 1987; Green & Bakus 1994), with the deepest record at 330 m in La Jolla, San Diego (Luke 1998). We therefore hypothesize that it is common from deep water up to ∼20 m and uncommon in shallower water. Not known from human-made structures.

## Discussion

The morphology and spiculation of *E. hyle* have been described in previous work (Aguilar-Camacho & Carballo 2013; Hooper *et al*. 1999). The freshly collected samples described here differ from previous descriptions in the nature of the ectosomal skeleton, which was previously described as vestigial, with thin styles tangential rather than in typical raspailiid brushes associated with protruding styles (Aguilar-Camacho & Carballo 2013; Hooper *et al*. 1999). We found typical raspailiid brushes, so this character is likely variable. Previous authors noted other variable traits as well, such as spicular dimensions and whether or not long styles are common (Aguilar-Camacho & Carballo 2013). The type location for this species is the Palos Verdes Peninsula, and one of the samples sequenced here (NHMLA 16545 / BULA-0541) is from this location. It would therefore be valuable to collect additional genetic data from Mexico and from deep-water samples to confirm conspecificity with the shallow-water *E. hyle* in California.

*Endectyon hyle* has been difficult for taxonomists to place. It was initially described as a *Hemectyon* (de Laubenfels 1932), which is now considered a subgenus of *Endectyon.* It was later assigned to both *Aulospongus* (Desqeyroux-Faundez & van Soest 1997) and *Raspailia (Raspaxilla)* (Hooper *et al*. 1999), based largely on the supposed presence of rhabdostyles. We find that nearly all acanthostyles in this species lack a bend near the head, and this is in fact consistent with previously published images of this spicule in this species (Hooper *et al*. 1999). In addition, de Laubenfels noted that the acanthostyles were located mainly in the periphery, which is now considered a trait of *Endectyon.* Moreover, the styles bear large, recurved spines, also a characteristic of *Endectyon*. Likely for these reasons, Lee et al. (2007a) included the species as a member of *Endectyon (Endectyon).* We agree that this placement is most consistent with the morphological characters. With additional support from the DNA phylogenies, we propose this species be placed in *Endectyon (Endectyon)*, pending a more systematic revision of the entire family.

Most samples of *E. hyle* found in California are probably identifiable in the field based on gross morphology alone. The most similar species is *Trikentrion helium*, which is also reddish-brown, stalked, and frondulose. Samples of *T. helium* differ by having large laminate fronds, each growing mostly in a single plane and lacking corrugations and wrinkles; *E. hyle* fronds are short, rugose and bushy, with multiple axes in a tangential section. However, the type specimen of *T. helium* (van Soest *et al*. 2012, fig. 24) appears somewhat bushy, likely because it was young and/or small, without extended blades. Further field work on the living morphology of these species is therefore warranted. Several small samples of *E. hyle* were also found that were short, rounded, and lumpy (figure 4B); these could be confused with *C. neon* in the field, though *C. neon* looks fleshier.

### Material examined

Holotype: CASIZ 236512, Coal Oil Point, Santa Barbara, California (34.40450, −119.87890), depth 3-6 m, 10/28/2019. Other samples: UCSB-IZC00041062 & SBMNH 693062, Cave Landings, San Luis Obispo, California (35.17535, −120.72240), intertidal, 2/6/2021; UCSB-IZC00041046, Arroyo Hondo, Santa Barbara, California (34.47182, − 120.14262), depth 4-8 m, 7/29/19; SBMNH 693087, Naples Reef, Santa Barbara, California (34.42212, −119.95154), depth 10-15 m, 7/31/2019; UCSB-IZC00041060, West End of Anacapa Island, California (34.01352, −119.44570), depth 7-12 m, 4/25/2019, collected by SP & TT; SBMNH 693084, Lechuza Point, Ventura, California (34.03437, −118.86146), intertidal, 12/14/2020; UCSB-IZC00041056 & SBMNH 693074, Goalpost Reef, San Diego, California (32.69438, −117.26860), depth 12-15 m, 2/8/2020; except where specified, all samples collected by TT.

### Etymology

From the latin *hispidus* (bristled) and *tumulus* (mound).

### Morphology

Thinly encrusting: the holotype is 5 mm thick, other samples 3-10 mm thick. Appearance usually evokes a rolling field of hills, like a barrow field; each hill is hispid, with smooth membranous depressions stretched between them. The thickest samples have deeper furrows between hispid mounds, such that they are more like hispid columns; in these cases, a cross-section through the sponge results in the sponge nearly falling apart into separate columns. Orange-red alive, like many other axinellid and microcionid sponges in the same habitats. Fades to cream/white in ethanol.

### Skeleton

Nodes of spongin ascend from a basal spongin mat, and both are sometimes cored with sediment. The basal mat contains a reticulation of smooth styles; this reticulation extends up from the mat in plumoreticulate columns bound with considerable spongin. Columns are echinated with smooth styles whose heads are embedded in spongin but are otherwise free; most echinating spicules are angled up giving the columns a columnar-cactus-like aspect. Plumoreticulated columns pierce the surface of the sponge to create hispid mounds; smaller and shorter reticulated columns are found between hispid mounds. Thin hair-like styles are embedded tangentially in the ectosome, singly or in bundles. Acanthostyles are present in the holotype but frequently absent in other samples. When present, they are found primarily with heads embedded in the basal spongin mat, tips up; some are also found echinating plumose columns.

### Spicules

Styles, acanthostyles, and thin ectosomal styles.

Styles: Generally curved or bent near head end; could perhaps be characterized as rhabdostyles, but this seems subjective. Size is variable among samples, with mean lengths varying from 254 μm to 403 μm (n = 20-49 per sample); mean width varies from 9 μm to 16 μm per sample. Holotype: 327-403-472 x 9-16-20 μm (n = 21). Combined distribution from all samples: 127-330-698 x 2-12-23 μm (n=271). One sample contained two oxeas.

Acanthostyles: Generally straight but sometimes curved. Unlikely to be characterized as rhabdostyles, as none show sharp bend near the head. Present in only 5 of 9 samples. Spines are often large and curved; most densely spined at tip end but occur all the way to the head of style. Mean size per sample from 151 to 171 μm in length, 9 to 12 μm in width; holotype 133-175-210 x 9-12-14 μm (n=21); combined distribution from all samples 118-166-224 x 6-11-20 μm (n=62). One sample contained a single acanthoxea.

Thin ectosomal styles: Very thin and usually broken; 133-196-332 x 1-3-4 μm (n=22).

### Distribution and habitat

Known range is from Avila Beach, Central California to Point Loma, Southern California, which is the Southernmost area investigated. A sponge photographed by Alison Young (California Academy of Sciences) at Yerba Buena Island, in San Francisco Bay, appears to be this species based on external morphology, so it may range into Northern California as well (inaturalist.org/observations/75598358).

This species is common in shady intertidal crevices, and is one of the most commonly encountered intertidal sponges in Southern California. Subtidally, it is common in very shallow water (<10 m), with a maximum collection depth of 15 m. It primarily grows on or under ledges in the subtidal, but is also found in the open, including in areas with high sedimentation where it can be found partially buried by sand.

## Discussion

This species should be placed in the subfamily Raspailiinae as it lacks the features characteristic of other raspailiid subfamilies. Placement in a genus is more difficult. Most encrusting species are placed in *Hymeraphia* (which can be excluded based on acanthostyle morphology) or *Eurypon*. *Eurypon* have a microcionid or hymedesmoiid skeleton, whereas *E. hispitumulus* is reticulated. The large curved spines on the acanthostyles are consistent with *Endectyon*, as is the close relationship with *E. hyle, E. delaubenfelsi,* and *E. fruticosum*.

Though the spiculation of this species varies among samples, there is no evidence to support subdivision into multiple taxa. Samples that lack acanthostyles do not differ from those that have them in terms of primary style size or other characters. Samples with DNA data include those with the longest and shortest styles, two with acanthostyles and two without, and samples spanning the Southern California portion of the range (Point Loma, Anacapa Island, Coal Oil Point, Arroyo Hondo); these DNA data contain no evidence for multiple species.

Though there are many orange/red encrusting sponges in California, the “hispid mound field” morphology often makes this species distinguishable in the field. This makes it all the more surprising that it has not been reported in any previous survey in California (Lee *et al*. 2007a). No described species nor undescribed morphospecies is similar to this species, despite considerable work on the Central California intertidal sponge fauna (Hartman 1975) and some surveys in Southern California (Bakus & Green 1987; Sim & Bakus 1986). It is therefore temping to speculate that this species is more common in this region today than in the recent past.

### Genus Raspailia (Nardo 1833)

Raspailia (Raspaxilla) hymani (Aguilar-Camacho & Carballo 2013; Hooper *et al*. 1999) Endectyon (Endectyon) hymani (Lee *et al*. 2007a) Aulospongus hymani (Desqeyroux-Faundez & van Soest 1997) Hemectyon hymani (Dickinson 1945; Green & Bakus 1994)

#### Material Examined

SBMNH 467982, Off Purisima Point, Santa Barbara (34.7757, −120.8367), 91-123 m depth, 7/23/1984, collected by Don Cadien.

#### Morphology

Growth mostly in a single plane, 13 x 11 mm. Surface a network of bundles 500-600 μm wide, forming a lattice-like pattern, fanning out towards distal end. Beige in ethanol, color alive unknown.

#### Skeleton

Choanosome contains a very dense reticulation of rhabdostyles, echinated by same. Long plain styles emerge from this axially compressed skeleton to pierce the surface of the sponge and extend up to 1700 μm past sponge surface. Thin styles also protruding from ectosome singly or in small groups, but characteristic raspailiid brush-like pattern not seen.

#### Spicules

Long styles, acanthorhabdostyles, and thin ectosomal styles.

Long styles: Gently curved or slightly sinuous, but not bent. 1333-1810-2157 x 26-27-30 μm (n=5).

Acanthorhabdostyles: Most have heads sharply bent at a 45-90 angle from shaft. Some have heads bent 180 degrees to form complete u-shaped head region, while a minority are straight. Spines are usually limited to the pointed half of the style but some are completely spined and some unspined. Size varies greatly in a continuous but bimodal distribution (modes at 175 and 305 μm). If separated into categories at 270 μm (following Aguilar-Camacho & Carballo (2013)), size distributions are 122-183-260 x 8-16-26 μm (n=59) and 272-313-384 x 21-27-32 μm (n=23); combined distribution 122-220-384 x 8-19-32 μm (n=82).

Thin ectosomal styles: Gently curved or slightly sinuous, but not bent. 498-601-967 x 3-5-9-6 μm (n=10).

#### Distribution and habitat

Only known from deep water, 90-138 m depth, from Purisima point (just North of the Southern California Bight) to Isla Partida, Cabo San Lucas, Mexico.

## Discussion

This specimen, previously examined by Green and Bakus (1994), is one of two specimens known from California. It differs from the published descriptions of two Mexican specimens in several respects including width of the lattice-like network of thickened sections (approximately 500-600 μm wide in California, 1200-2000 μm wide in Mexico based on figure 1 of Aguilar-Camacho & Carballo (2013)) and the length of the thin ectosomal styles. Most notably, the very long protruding styles that are prominent in the California specimen are not mentioned and presumed absent in the Mexican samples. This difference is similar to the variation reported above in *E. hyle*, and the similarities between the California and Mexican material are notable. All samples are fan shaped with a lattice-like pattern of ridges, and have acanthostyles that are severely bent at the head. Splitting the size distribution of acanthosyles in the same way as Aguilar-Camacho and Carballo (2013) leads to very similar size distributions in both size bins. We therefore think it is likely that the California and Mexican samples are of the same species. DNA extractions were attempted but did not yield PCR amplifiable DNA.

### Genus Eurypon (Gray 1867)

#### Material examined

Holotype CASIZ 236513; other samples SBMNH 693059, UCSB-IZC00041061; all collected by TT at Bird Rock, San Diego, California, USA (32.81447, −117.274313), in the intertidal zone, 1/10/2021.

Etymology

The distinctly curved acanthostyle resembles a bent (*curvus*) nail (*clavus*).

#### Morphology

Thinly encrusting. Holotype approximately 6 x 6 cm, 1 mm thick; other samples 2-6 cm across, 0.5 −1 mm thick. Surface fairly flat, not particularly rugose, but varying thickness makes the surface slightly undulating. Covered in small pores of varying size, from about 20-100 μm in diameter. Visibly hispid due to long protruding styles. Reddish orange when alive, beige when preserved in ethanol.

#### Skeleton

Sponge contains a basal plate of spongin with many acanthostyles embedded within. Vertical trunks of spongin arise from the basal plate and are cored with styles; some styles have tips free, making columns plumose. Columns are echinated with acanthostyles and sometimes bridged by horizontal tracks of spongin cored with styles. Long thin styles are embedded near the tops of some columns; these pierce the surface of the sponge and make the surface hispid. Thin ectosomal styles fan out in brushes around these protruding styles, creating the typical raspailiid ectosomal skeleton.

#### Spicules

Styles, long thin subectosomal styles, acanthostyles, and thin ectosomal styles.

Styles: straight, slightly curved, or weakly bent; if bent, bend is near head end, unlike acanthostyles. Average length varies from 313 to 350 μm; holotype: 238-350-520 x 5-10-13 μm (n=23); all samples pooled: 193-338-633 x 5-10-14 μm (n=55). Long thin subectosomal styles: straight or gently curved; sometimes with weak subterminal swelling near head. Generally longer and thinner than coring styles, but size classes are hard to clearly separate and likely overlap. Dimensions in holotype: 550-676-945 x 3-5-6 μm (n=8); all samples pooled: 491-760-994 x 3-5-8 μm (n=21).

Echinating acanthostyles: most are sharply bent; in contrast to rhabdostyles, bend is usually between 50% and 66% of the way towards tip. Most are weakly spined near tip only, but some are completely unspined; rarely (<1%) modified to oxeas. When measured in a straight line from head to tip, dimensions in holotype are 106-135-166 x 4-6-10 μm (n=33); all samples pooled: 84-127-215 x 3-6-12 μm (n=99); average lengths for each sponge individually = 112, 133, 135 μm. If lengths are measured along the curve, lengths increase approximately 10%.

Thin ectosomal styles: only 1-2 microns thick, lengths from 100-500 μm, but most were broken and likely longer in life.

#### Distribution and habitat

All samples were found on short rock walls or under ledges, exposed at low tide, at Bird Rock in San Diego. Three samples were found in less than an hour of searching, so it seems to be abundant at this location. As it was not reported in previous surveys in California (Bakus & Green 1987; Hartman 1975; de Laubenfels 1932), nor found at any other location in the current work, nor reported from surveys in Mexico (Carballo *et al*. 2019; Gómez *et al*. 2002; Hofknecht 1978), it doesn’t seem to be common at many other locations.

## Discussion

The ectosomal skeleton, echinating spicules, and lack of polyactines place this species in the subfamily Raspailiinae. The encrusting morphology of this sponge, with its “microcionid” skeletal structure (fiber nodes ascending from a basal layer of spongin) are most consistent with placement in the genus *Eurypon*. *Eurypon* are defined as “typically encrusting Raspailiidae with microcionid skeletal structure in which fiber nodes ascend from the basal layer of spongin” (Hooper 2002c). Many of the described species lack these ascending fiber nodes, and instead have a “hymedesmoid” skeletal structure of spicules embedded directly in the basal spongin, tips up (Aguilar-Camacho & Carballo 2013; Cavalcanti *et al*. 2018; Santos *et al*. 2014). This variable skeletal structure may be related to the thickness of the species (Santos *et al*. 2014). Previous molecular phylogenies of multiple loci found *Eurypon* to be polyphyletic (Redmond *et al*. 2013; Thacker *et al*. 2013), which is also found in our updated phylogenies. A revision of the genus awaits genetic data from more species and a better resolved phylogenetic tree.

The bent echinating styles differentiate this species from all described species of *Eurypon*, as listed by Recinos et al. (2020). There are also four species of encrusting Raspailiinae described in the genus *Hymeraphia*, though none are known from the Pacific; all *Hymeraphia* greatly differ from the newly described species in echinating style morphology and skeletal architecture (Morrow *et al*. 2018).

No other Eurypon have been described from California; the closest records are several *Eurypon* with a more hymedesmoid architecture known from the tropical Mexican Pacific (Aguilar-Camacho & Carballo 2013). A “*Eurypon? sp.*” was also listed as occurring in British Columbia, but with no published data on its morphology (Austin 1985).

Identifying this species in the field would be challenging, as there are many thinly encrusting intertidal Microcionidae in California; the smooth, non-rugose surface of this sponge is very similar to sympatric *Clathria* such as *C. originalis*. These *Clathria* lack the long protruding styles, which are visible in good macro photos of live colonies; *Cyamon koltuni* are not known from the intertidal, but are also hispid with protruding styles.

### Genus Aulospongus (Norman 1878)

### Material examined

Holotype: CASIZ 236515 / UCSB-IZC00041063, Big Rock, Santa Cruz Island, California, USA (34.05220, −119.57360), 6-14 m depth, 1/19/2020.

#### Etymology

From the latin *viridis*, meaning green.

#### Morphology

Massive and irregular, approximately 5 x 3 x 2 cm. Surface covered in prominent conules, with oscula concentrated at high points. Dark green in life, it retains its pigmentation in ethanol. Surface membrane is opaque after preservation, but in life it is a mix of opaque regions and regions with a visible mesh of pores.

#### Skeleton

Choanosome contains plumose ascending tracts of spongin cored with styles, 50 − 250 μm thick, becoming less dense at the periphery. Tracts are echinated by spined and unspined styles. Coring styles tend to be longer, straighter, less tylote and less spined, on average, but this is not absolute. Long subectosomal styles pierce the ectosome and thin ectosomal oxeas form brushes that pierce the surface.

#### Spicules

Rhabdostyles, long subectosomal styles, and thin ectosomal styles.

Rhabdostyles: usually with a terminal tyle, but sometimes with a subterminal bulge, polytylote, or plain. Spines are small when present, and usually limited to the pointed half of the style, but many are entirely smooth. Spined and unspined rhabdostyles are difficult to separate, as some are very weakly spined; when pooled, the length distribution is bimodal, with modes at 215 and 315 μm. Acanthorhabdostyles alone: 103-249-420 x 3-11-17 μm (n=109); unspined rhabdostyles alone: 156-300-528 x 5-10-14 μm (n=30); Wilcox rank-sum test indicates acanthorhabdostyles may be smaller (p=0.06).

Subectosomal styles: sometimes curved or bent but generally lacking sharp bend of rhabdostyles and most lack tyle; 565-839-1091 x 5-10-15 μm (n=12). Ectosomal styles/strongyloxeas: Thin ectosomal styles most common, but some are fusiform and some pointed at both ends; 246-456-679 x 2-3-5 μm (n=34).

#### Distribution and habitat

The holotype is the only sample found to date. Found at Big Rock on Santa Cruz Island, which is a sheltered cove with shallow rocky reef and kelp forest. The sponge was found in a shaded crevice among large boulders.

## Discussion

Raspailiidae with rhabdostyles are currently allocated to three taxa: *Aulospongus*, *Raspailia (Raspaxilla)*, and *Endectyon (Hemectyon)*. In *Raspaxilla* and *Hemectyon*, fibres are cored by smooth, non-rhabdose styles (Hooper 2002c). We find the distinction between rhabdostyle and non-rhabdose bent or curved styles to be subjective, and more of a spectrum than distinguishable categories. In the case of *A. viridans*, however, the distinctly asymmetric tyle found on both coring and echinating styles seems likely to fall on the rhabdostyle end of the spectrum, and it is found in large unspined styles and smaller ones that are entirely spined. This sponge also lacks distinction between axial and extra-axial skeleton, and the plumose ascending tracts are more dendritic than reticulated, and both these traits are also more consistent with *Aulospongus* than *Raspaxilla* or *Hemectyon* (Hooper *et al*. 1999). The green color of this sponge, and the fact that it retains its pigmentation in ethanol, are in contrast to all other sponges described herein. This color also sets it apart from all known species of *Aulospongus* (and *Raspaxilla*). Three species of *Aulospongus* have been described from the Pacific, all from Mexico. Two, *A. cerebella* (Dickinson 1945) and *A. californianus* (Aguilar-Camacho & Carballo 2013), are deep sea species, pale beige in ethanol, tubular or vase-shaped, and have much thicker styles. The final Pacific species, *A. aurantiacus* (Aguilar-Camacho & Carballo 2013), is known from shallow water but is orange, encrusting, and lacks a raspailiid ectosomal skeleton.

All photos from holotype.

### Material examined

Holotype: P119, Hopkin’s Bridge, 17 m West of Quast Rock, Point La Jolla, California, USA, 21-25 m depth, 8/10/1965.

### Etymology

Named for La Jolla, California.

### Morphology

Holotype is three fragments, up to 44 mm thick and 67 mm across. Color alive unknown; brown preserved. Hispid and extremely rugose, the sponge surface is covered in tufts and fan-like ridges several millimeters in height.

### Skeleton

Choanosome contains plumose ascending tracts of spongin cored with, and echinated by, acanthorhabdostyles; tracts branch dendritically. Tracts become less dense and thinner towards the periphery. Long subectosomal styles pierce the ectosome, surrounded by thin ectosomal oxeas in disorganized clumps and bundles. Acanthorhabdostyles also pierce sponge surface when tracts run parallel to it.

### Spicules

Acanthorhabdostyles, long subectosomal styles, thin ectosomal oxeas/strongyloxeas/styles.

Acanthorhabdostyles: bent near head, or nearly straight but with tyle placed asymmetrically to one side. Spines usually small, variable in extent but average coverage is about half of style, usually limited to pointed end; spined rarely occur to head region; less than 5% are unspined. Length varies considerably but size distribution is unimodal. Rare aberrations included one acanthoxea, spined on only half its length, and one spined quadractine. 185-333-537 x 6-15-26 μm (n=101).

Long subectosomal styles: Smoothly bent or slightly sinuous, with very weakly tylote heads. 1497-1903-2304 x 9-15-21 μm (n=17).

Ectosomal oxeas/strongyloxeas/styles: Some clearly oxeas, with two pointed ends, others are nearly styles, but with head end slightly tapering; intermediates also found. 494-695-818 x 3-5-9 μm (n=18).

### Distribution and habitat

The holotype is the only known sample. It was collected at 21-25 m depth near La Jolla, in extreme Southern California.

## Discussion

Like *A. viridans*, this species has a distinctly asymmetric tyle on the acanthostyles, plumose ascending tracts that are more dendritic than reticulated, and a lack of distinction between axial and extra-axial skeleton. These traits are most consistent with *Aulospongus* (Hooper *et al*. 1999). This species differs from *A. viridans* in color, growth form, fewer unspined rhabdostyles, and spicule dimensions (all spicule categories are much longer in *A. lajollaensis*). Indeed, the styles of this species are twice as long as any described *Aulospongus*, and the oxeas are longer than all species except *A. involutus* (Kirkpatrick 1903), a species from the Indian Ocean (Cavalcanti *et al*., 2014). Additional traits differentiate it from species known from the Eastern Pacific: *A. californianus* has two sizes of acanthorhabdostyles, while *A. lajollensis* has a broad but unimodal size distribution; *A. auranticus* has entirely spined acanthostyles and a different growth form; *A. cerebella* lacks oxeas and is not hispid (Hooper *et al.,* 1999).

### Subfamily Cyamoninae (Hooper 2002)

### Genus Trikentrion (Ehlers 1870)

Trikentrion helium (Claisse *et al*. 2018; Dickinson 1945; Smith 1968; van Soest *et al*. 2012) Trikentrion catalina (Gómez *et al*. 2002; van Soest *et al*. 2012) Cyamon catalina (Lee *et al*. 2007a; Sim & Bakus 1986)

#### Material examined

Collected by TT: NHMLA 20362, Hawthorne Reef, Los Angeles, California, USA (33.74714, −118.42090), 24 m depth, 8/23/2019; UCSB-IZC00041058,

Trainwheels Reef, San Diego, California, USA (32.65205, −117.26243) 28 m depth, 9/19/2020. Collected by others: USNM 33631 (*Trikentrion catalina* holotype), Bird Rock, Catalina Island (33.45, −118.40), 50 m, date not recorded; NHMLA L35535 (*Trikentrion helium* holotype), South Bay, Pacific Mexico, (28.07, −115.3), 18–27 m, 3/10/1934.

#### Morphology

One or more thin bladed lamina (2-3 mm thick) growing from a stalk. Each leaf-like blade has growth mostly in a single plane, with even thickness, lacking corrugations. Very hispid. Consistency firm and leathery. Bright red to orange-red alive, beige in ethanol.

#### Skeleton

A dense mass of polyactines cores each blade, surrounded by tangential styles. Long styles are perpendicular to the core, protruding from the ectosome, surrounded by typical raspialiid brushes of smaller styles.

#### Spicules

Long thin styles, short thin styles, polyactines, trichodragmas. Measurements shown below are from both newly collected samples combined.

Long thin styles: 1135-1709-2708 x 6-13-22 μm (n=22).

Short thin styles, usually plain styles but sometimes few tylostyles; some fusiform, approaching strongyloxeas/oxeas: 364-527-673 x 2-4-8 μm (n=24).

Polyactines: predominantly triactines with one shorter (basal) cladus which usually bears prominent spines and two lateral clades which are unspined. Shape often sagittal, with unspined clades at a 180-degree angle. Variation is considerable, with many triactines at various angles, sometimes entirely unspined, and with length of clades deviating from the typical pattern. Occasional tetractinal polyactines. Diactinal polyactines numerous and diverse in shape. Many have one cladus replaced with a knob, while others have two clades. The two clades may be at an angle or straight; when straight the shape is like an oxea, but with a swollen region where the clades connect. One ray of the diactines is sometimes spined. Basal clades of triacts: 51-75-110 x 8-15-20 μm (n=35), lateral clades of triacts: 57-128-200 x 5-13-20 μm (n=32). Straight diacts 133-210-311 μm in total length, 9-13-22 μm wide at widest point excluding enlarged central knob (n=9).

Trichodragmas: 67-75-88 x 7-12-18 μm (n=13); width of individual raphides not measured.

#### Distribution and habitat

Rocky reef. Depth range 17-94 m. Known from Palos Verdes, Los Angeles County, California to Guerrero, Mexico. We found it to be common, but not abundant, on reefs below 20 m in Los Angeles and San Diego Counties.

## Discussion

*Trikentrion helium* was described from Cedros Island, Baja California, Mexico (Dickinson 1945). It was later collected at various locations around San Diego, California (Luke 1998; Smith 1968), and these collections were used in the chemical analyses that characterized the structure of the carotenoid now known as trikentriorhodin (Aguilar-Martinez & Liaaen-Jensen 1974). More recently, it was reported as common at reefs around the Palos Verdes Peninsula, Los Angeles, California (Claisse *et al*. 2018).

*Trikentrion catalina* was described from two sponges collected at Catalina Island, Southern California (Sim & Bakus 1986). Five sponges from the Mexican tropical Pacific, from Nayarit and Guerrero, were later ascribed to this species as well (Gómez *et al*. 2002). The authors of both papers on *T. catalina*, however, appeared to have overlooked *T. helium*, as this species is extremely similar but was not mentioned. Van Soest et al. (2012) re-described the holotypes of both species and verified that they were closely related, noting only two differences. First, the *T. helium* holotype contained diactinal polyactines and the *T. catalina* holotype did not. Second, growth form was subtly different: “undulating thin-bladed sheets together forming a bushy mass” in *T. helium* vs. a single stalked blade growing mostly in a single plane in *T. catalina*. The freshly collected samples described here have diactinal polyactines and would therefore be ascribed to *T. helium*, but the growth forms are intermediate, with multiple hin fronds emerging from a base. The fresh sample from the Palos Verdes peninsula was collected 30 km from the type location for *T. catalina*, and these locations are biogeographically similar.

We therefore suspected that *T. catalina* was a junior synonym of *T. helium*. To investigate further, we rexamined the holotypes of each species. We were able to find numerous diactines in the holotype of *T. catalina*. Diactines that are completely straight (figure 11A, B) were quite rare, with only one found on the existing slide previously examined by Van Soest *et al*. Searching through a large number of spicules isolated from another fragment of the holotype yielded additional examples, as well as a larger number of bent diactines (figure 11C) and a large number of transitional forms with one clade replaced by a knob (figure 11D). All polyactine forms seen in *T. helium* were located in the *T. catalina* holotype. As this leaves no morphological character to distinguish these species, *T. catalina* should be considered a junior synonym of *T. helium*. *Trikentrion helium* is distinctive and should usually be identifiable in the field based on gross morphology alone. The most similar known species is *E. hyle*; see discussion section above for that species for additional information.

### Genus Cyamon (Gray 1867)

Cyamon neon (Lee *et al*. 2007a; Sim & Bakus 1986; Smith 1968; van Soest *et al*. 2012)

### Material examined

Samples collected by TT: NHMLA 16556 / BULA-0546, Hawthorne Reef, Palos Verdes Peninsula, Los Angeles (33.74714, −118.42090), 18-24 m, 8/23/2019; UCSB-IZC00041065 and TLT11, Elwood Reef, Santa Barbara (34.41775, −119.90150), 9-14 m, 4/17/2019; SBMNH 693065, Elwood Reef, Santa Barbara (34.41775, −119.90150), 9-14 m, 4/26/2019; SBMNH 693055, Arroyo Quemado Reef, Santa Barbara (34.46775, −120.11905), 7-11 m, 7/29/2019; SBMNH 693056, Arroyo Quemado Reef, Santa Barbara (34.46775, −120.11905), 7-11 m, 6/14/2019; SBMNH 693103, UCSB-IZC00041052 and UCSB-IZC00041053, Parson’s Landing, Catalina Island (33.47502, −118.55000), 8-17 m, 11/10/2019; Samples collected by others: NHMLA 15920 / BULA-0229, Marineland, Palos Verdes, Los Angeles (33.75166667, −118.41665), depth not recorded, collected by Gustav Paulay; NHMLA L35535 (*Cyamon argon* holotype), South Bay, Cedros Island, Mexico, 18-27 m, 3/10/1934; USNM 21384 (*Cyamon neon* paratype), South of San Pedro, Los Angeles, 36 m. 9/24/1924; P109, Quast Rock, 2 km WNW of Point La Jolla, San Diego, (32.75, −117.37), 23 m, 5/1/1965; P114, San Onofre Power Station, 0.8 km off South pier, San Diego, (33.368, −117.555), 15 m, 8/19/1965; P111, South Cash Beach, 0.8 mi out, La Jolla, San Diego, (32.75, −117.37), 11 m, 8/6/1965; P113, collection data lost, but likely collected near La Jolla, San Diego in 1965, based on collection data from numbered samples before and after.

#### Morphology

Thickly encrusting (1-2 cm) or with upright lobes up to 5 cm tall; larger sponges become a bushy mass. Internal columns of orange flesh end in hispid conules, making sponge surface extremely rugose. Covered in a whitish, partially transparent, porous dermal membrane; large pores (approx. 0.5 mm) in this membrane create a spider-web-like aspect when sponge is alive and relaxed. Oscules occur singly or in pairs, primarily in elevated areas, each 1-2 mm in diameter. Orange in life; reacts to ethanol and turns very dark brown.

#### Skeleton

Thick (200-400 μm wide) columns of spongin, styles, and polyactines ascend towards the sponge surface, splitting into thinner columns towards the ectosome. Thinner (< 100 μm) columns of spongin cored with styles form sparser, plumoreticulated regions beneath conules, with fewer polyactines present. Centrylote strongyles are found in tangential bundles near the ectosome, but also scattered throughout.

#### Spicules

Short styles, long styles, wavy centrylote strongyles, polyactines (with 2-5 clades).

Short styles: curved or slightly bent near head end; sometimes fusiform due to constriction near head region; rarely modified to oxeas or strongyles; 251-442-645 x 3-18-40 μm (n=438).

Averages per sponge vary from 383 to 539 μm in length and 11 to 29 μm in width (see discussion below).

Long styles: curved or slightly bent near head end; 855-1306-2264 x 4-12-25 μm (n=59).

Wavy centrylote strongyles: thin with blunt ends (strongylote) or with one or both ends pointed (style/oxea); often but not always with a centrylote swelling; 177-296-482 x 2-5-8 μm (n=27). Diactinal polyactines: unspined, with irregular centrylote swelling; straight or bent at the centrylote swelling. 58-200-373 x 3-14-29 μm (n=368). Averages per sponge vary from 172 to 234 μm in length and 8 to 22 μm in width.

Other polyactines: mostly three claded, but occasionally with 4 or 5 clades. Highly variable angles between clades, with some equilateral, some sagittal, and some with the widest angle more than 180 degrees. One (basal) clade often shorter than other two. Clades end in rounded, blunt tips or sharp points. Entirely unspined or with small spines or bumps present on basal clade; rarely spined on multiple clades. Basal clades 24-50-78 x 6-14-24 μm (n=251); lateral clades 42-97-169 x 5-14-26 μm (n=293).

#### Distribution and habitat

Known from Southern California to Cedros Island, Baja California, Mexico. Depth range 7-36 m (though Lee et al. (2007a) report that it is intertidal, we know of no published reports from the intertidal, nor did we find any intertidally). We found it to be common but patchy in Southern California, locally abundant in some places but absent from others. In total, it was found at 11 of 52 natural rocky reefs surveyed in Southern California, without any apparent geographic pattern in abundance. It was not found on any human structures (wrecks, pilings, floating docks, or oil rigs).

## Discussion

Cyamon neon was described from three samples collected near San Pedro in Southern California (de Laubenfels 1932). de Laubenfels noted that the sponge was dark brown, and that the ectosome was full of dark brown granules, which we can now report are formed when the sponge reacts to ethanol.

A very similar species, *Cyamon argon* (Dickinson 1945), was later described from a single sample collected at Cedros Island, Baja California, Mexico. The holotypes were compared by van Soest et al. (2012), who noted their similarity but retained both species. The differences originally noted in the *C. argon* description were revised by later authors, but three differences were noted by von Soest et al., as follows. First, the short styles and diactines were considerably thicker in the *C. argon* holotype than the *C. neon* holotype. Second, the basal clades of the polyactines are more prominently spined in the *C. argon* holotype than the *C. neon* holotype.

Finally, the *C. neon* holotype is thickly encrusting while the *C. argon* holotype is more upright and bush-like. In his PhD thesis, however, Smith (1968) examined the a larger sample of material: holotypes, paratypes, and several more samples from La Jolla California. The paratype of *C. neon* was much like the holotype of *C. argon,* and the La Jolla samples were intermediate, leading Smith to conclude that the differences between holotypes were part of a continuous spectrum of variation. He therefore proposed *C. argon* was a junior synonym.

To further investigate the distinctiveness of these named taxa, we reexamined the holotype of *C. argon*, the paratype of *C. neon,* the La Jolla samples of Smith, and seven freshly collected samples. DNA sequencing was only successful on fresh samples, but our morphological data agree with Smith: we find that all differences between holotypes are part of a spectrum of continuous variation, and that the *C. neon* paratype is very like the *C. argon* holotype. Figure 12 shows the spicule widths of diactines (A) and styles (B). The difference between holotypes is substantial, but the paratype of *C. neon* has the thickest spicules of any sample, and the other samples show that variation is continuous. The spination of triactines was likewise variable. Though variation was continuous, we attempted to quantify spination by classifying samples as having mostly spined basal clades, a mix of spined and unspined basal clades, and basal clades that were only rarely spined (black, gray, and white respectively in figure 12A). The *C. neon* paratype had the highest proportion of spined basal clades, with some La Jolla samples intermediate, and other samples largely unspined. Finally, with respect to growth form: we found samples with *C. neon* like spicule widths but with vertical growth, and it was previously reported that *C. neon* in deeper waters are more erect and branching, with heights up to 10 cm (Smith 1968). As shown in figure 11C, *Cyamon* from the type locality for *C. neon* grow to be very bush-like at depths below 20 m.

**Figure 12.**
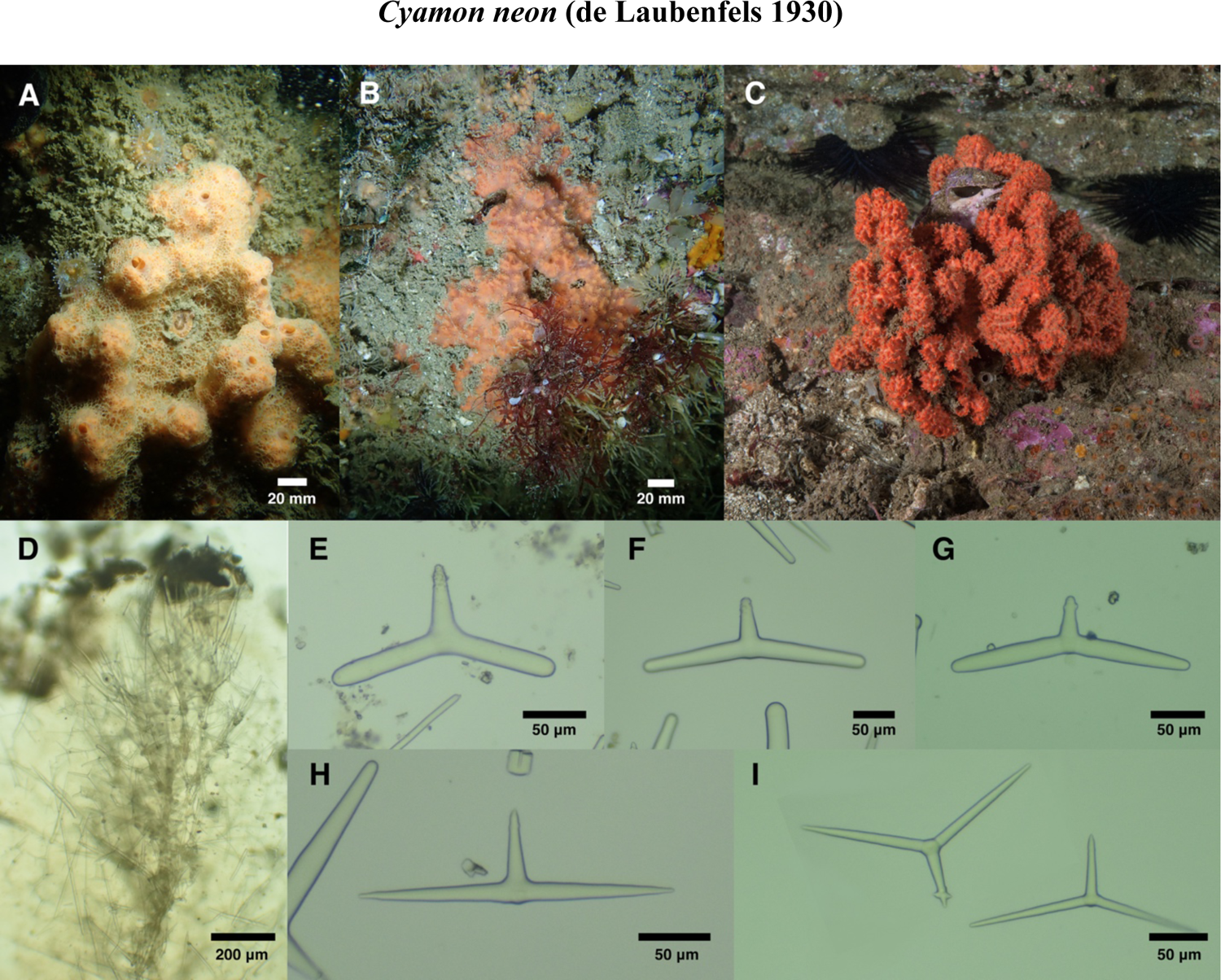
*Cyamon neon.* Field photos of A: TLT11, B: UCSB-IZC00041053, and C: an uncollected bush-like sample, photographed by Mary Passage at Resort Point Reef, off the Palos Verdes Peninsula, at approximately 21 m depth. D: Skeletal structure of SBMNH 693055 at the periphery. E: Variety of diactinal polyactines from UCSB-IZC00041052. F: Polyactines from UCSB-IZC00041052. G: Wavy centrylote strongyle/style from UCSB-IZC00041052.

**Figure 13.**
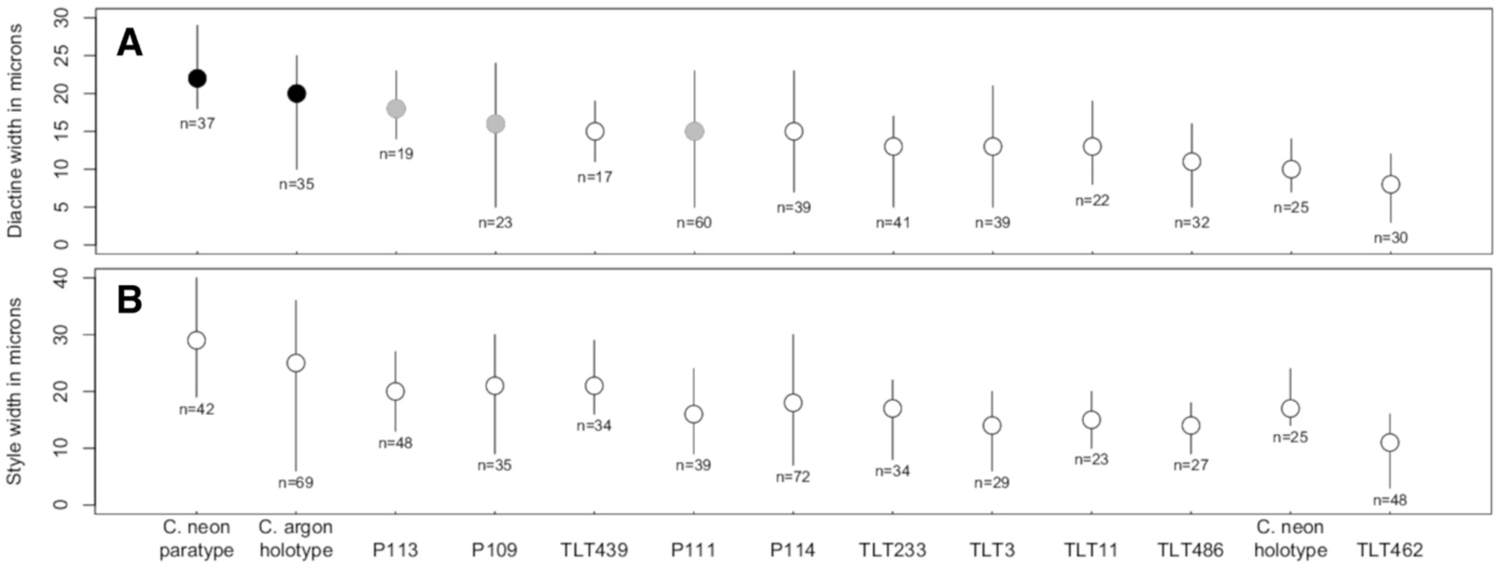
Distribution of spicule widths in *Cyamon neon.* Mean values (circles) and range (lines) are shown, with sample size indicated for each sponge. A: diactines; shading indicates the proportion of triactines with spined basal clades (black=most, gray= many, white=few). B: styles. Data from the *C. neon* holotype are from Van Soest *et al*. (2012); all other data collected by authors.

**Figure 14.**
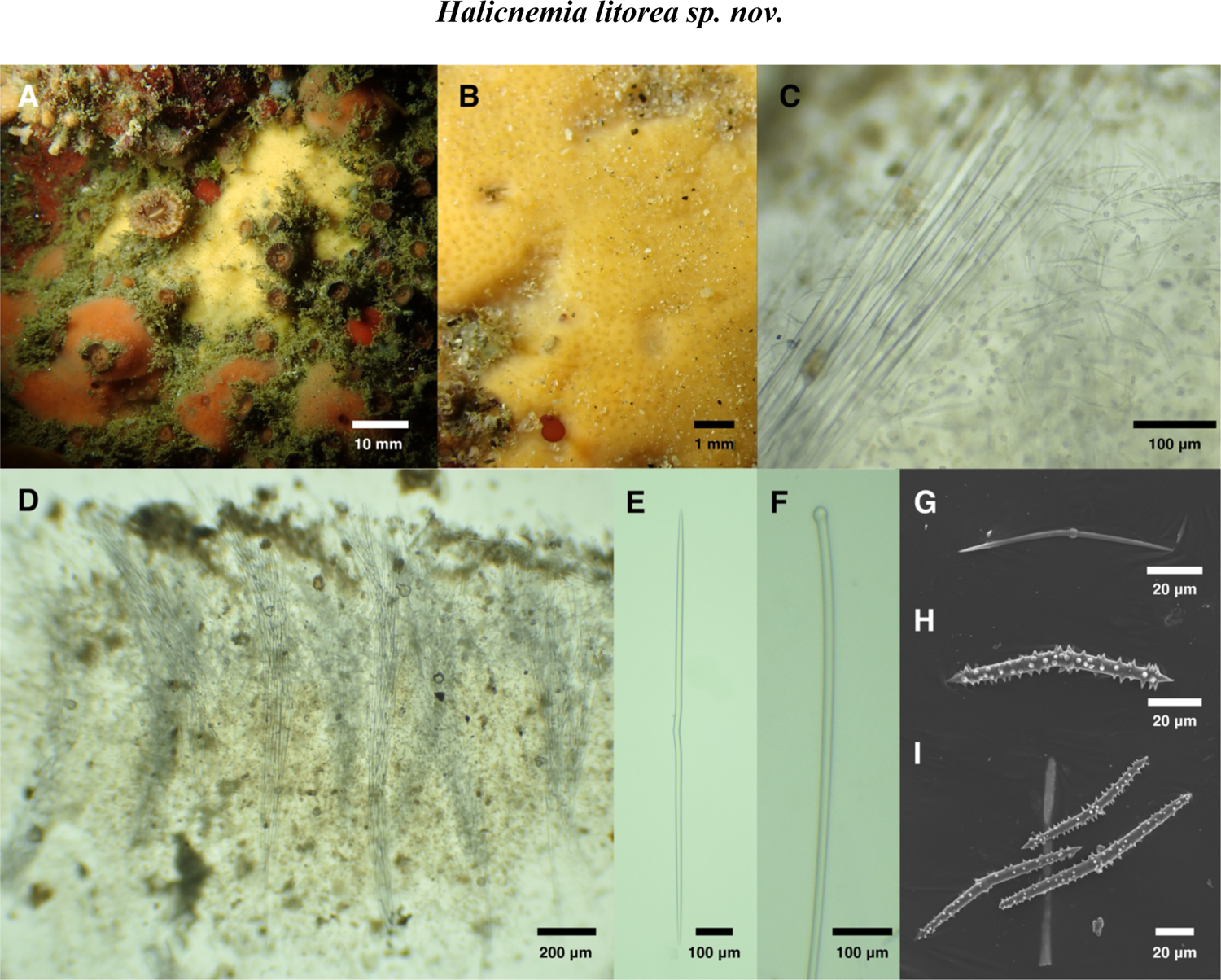
*Halicnemia litorea.* A: Field photo of holotype (CASIZ 236514). B: Field photo of sample SBMNH 693060. C: Skeletal structure of holotype. D: Skeletal structure of sample UCSB-IZC00041045. E-I: Spicules from holotype. E: Oxea with central kink, F: Head of tylostyle, G: centrylote microxea, H-I: Acanthoxeas displaying variable spination.

**Figure 15.**
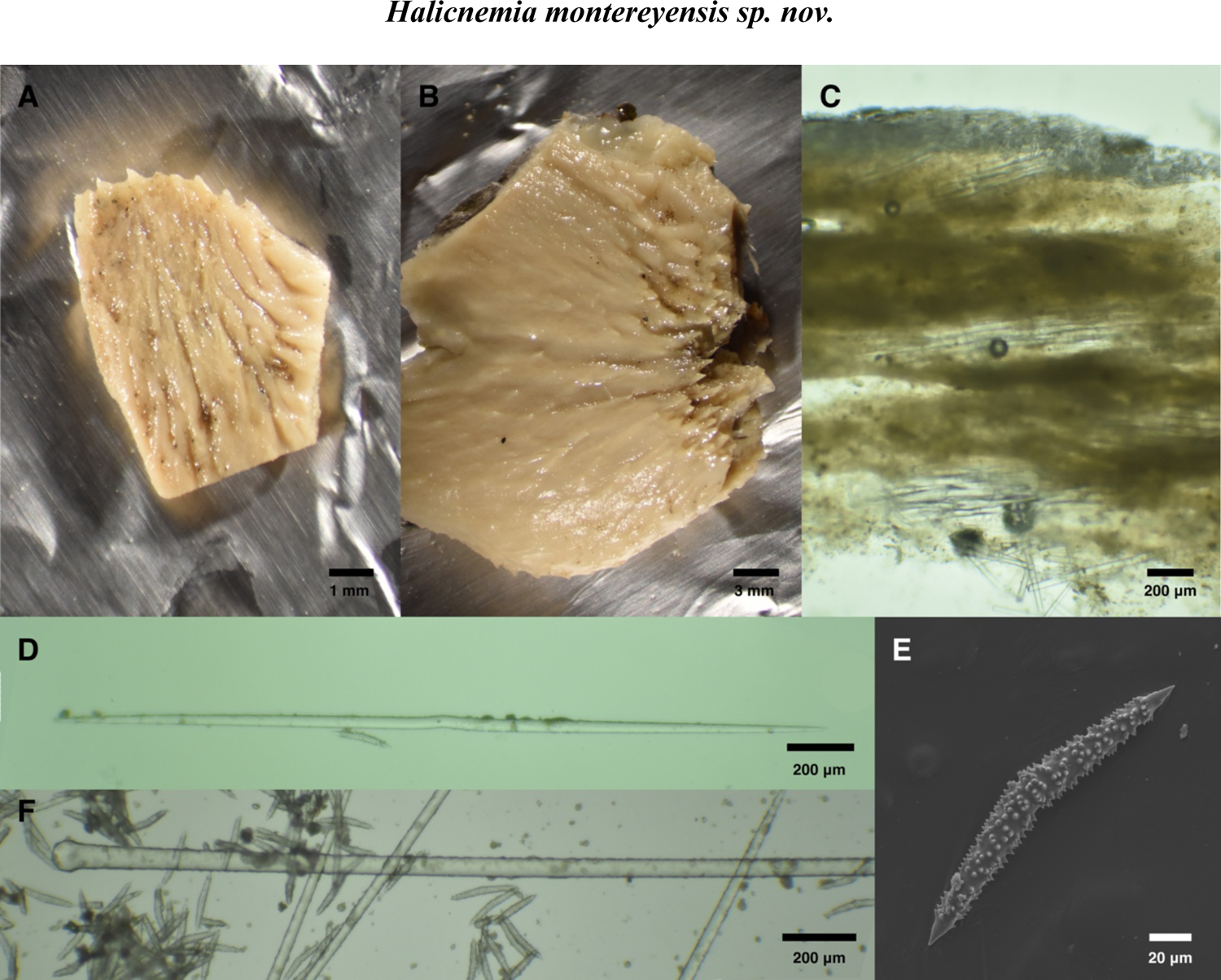
*Halicnemia montereyensis*. A: Holotype, CASIZ 165841, post-preservation. B: CASIZ 165844, post-preservation. C: Tangential section of holotype showing spicule bundles and crust of acanthoxeas. D: Kinked oxea from CASIZ 165844. E: Acanthoxea from CASIZ 165838. F: Head of tylostyle from CASIZ 165838.

**Figure 16.**
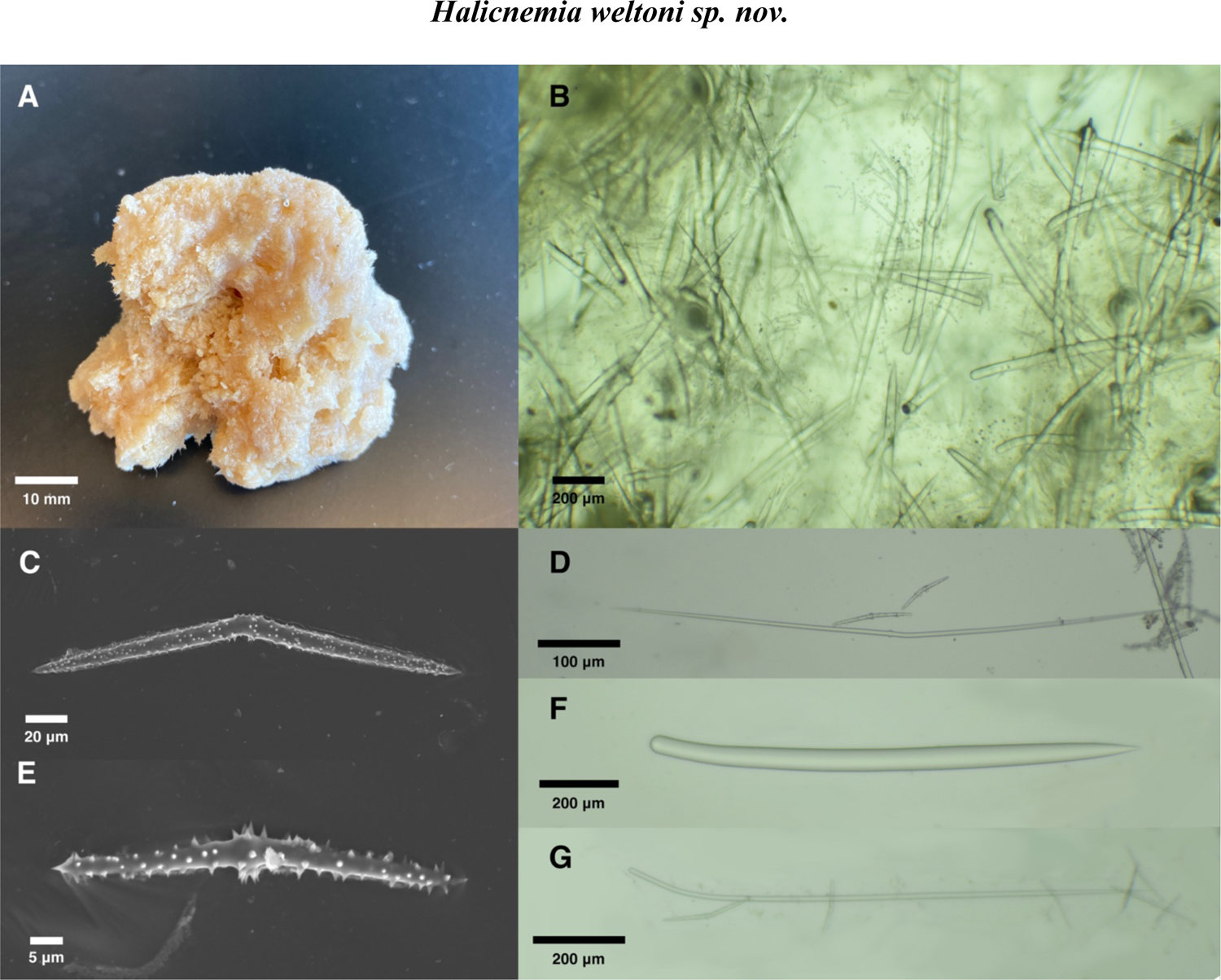
*Halicnemia weltoni.* A: Holotype, CASIZ 108411, post-preservation. B: Tangential section of CASIZ 184380. C, E: Acanthoxeas from holotype. D: Centrylote oxea from CASIZ 184380. F: Thick style from holotype. G: Thin style from holotype.

These results clearly indicate that there are no morphological traits that distinguish *C. argon* from *C. neon*. There are also no notable habitat differences, as the *C. argon* holotype was collected on shallow (18-27 m) rocky reef among giant kelp, which is the same habitat where the *C. neon* samples have been found. *Cyamon argon* should therefore be considered a junior synonym of *C. neon*.

*Cyamon koltuni* (Sim & Bakus 1986)

### Material examined

USNM 33630 (holotype), Big Fisherman’s Cove, Catalina Island (33.45, −118.48), 6m, date not recorded, spicule slide only.

#### Morphology

Thinly encrusting, up to 1 mm thick. Bright orange and hispid.

#### Skeleton

Basal mass of polyactines with emergent styles.

#### Spicules

Long thin styles, short thin styles, short thick styles, and polyactines. For dimensions see van Soest et al. (2012).

#### Distribution and habitat

Known from two samples, one from Big Fisherman’s Cove, Catalina Island, California, and the other from the Islas Marietas, Nayarit, Mexico.

## Discussion

The holotype is the only known sample from California. We examined the previously prepared slide of spicules from the holotype, but found few spicules present on the slide. However, an second sponge of this species was found in Pacific Mexico, as described by van Soest et al. (2012). This species is easily distinguished from other California species by possession of polyactine spicules with bulbous spined tips. It is unlikely to be distinguishable in the field due to the many thinly encrusting red and orange sponges known in California.

**Family Stelligeridae (von Lendenfeld 1898)**

**Genus Halicnemia (Bowerbank 1864)**

*Higginsia* ?*higginissima* (Ristau 1977)

*Higginsia sp.* Bakus and Abbot 1980 (Bakus & Abbott 1980; Klontz 1989)

*“Higginsia* sp. Klontz, 1989”, in part (Lee *et al*. 2007a)

### Material examined

Holotype: CASIZ 236514, Otter Cove, Monterey, (36.62920, −121.92031), depth 5-12 m, 11/24/2019, collected by TT; UCSB-IZC00041045, Isla Vista Reef, Santa Barbara, (34.40278, −119.85755), depth 5-12 m, 8/1/2019, collected by TT; UCSB-IZC00041047, Coal Oil Point, Santa Barbara, (34.40450, −119.87890), on loose stone in a tidepool, 7/6/2019, collected by TT; SBMNH 693060, Cannery Row, Monterey, (36.61798, −121.8978), depth 9-16 m, 9/21/2021, collected by TT; CASIZ 66860, SE Farallon Island, “mussel bed study site and Jewel Cave”, intertidal, 12/6/1975, collected by “Chaffee, Bowman, and Heinonen” (Farallon Research Group); CASIZ 108558, Santa Rosa Island, “1.2 mi E of Fort Point”, 12-14 m depth, 8/29/1985, collected by Sarah Ward Klontz; CASIZ 178096, Pigeon Point, San Mateo County, intertidal, 7/17/2004, collected by “Moss Landing Marine Lab”; CASIZ 180207, Channel Islands Harbor, Ventura, 0-3 m depth, 7/25/2006, collected by “Moss Landing Marine Lab”.

#### Etymology

Named for its habitat (the littoral), which is unusual for the genus.

#### Morphology

Thinly encrusting. Holotype was approximately 4 x 4 cm in life and 2 mm thick; other samples 2-4 mm thick. The four freshly collected samples, including the holotype, were yellow when alive; where noted in museum records, other samples were yellow, yellow-green, yellow-orange, or orange at c eserved samples are beige to pale yellow. Several samples, including the holotype, were notably slimy at collection; slime was not noted for the other samples but may have been present. Hispidity is apparent in field photos but difficult to see with the naked eye.

#### Skeleton

Bundles of megascleres rise vertically from the substrate, fanning out slightly as they approach and pierce the surface to create hispidity. Acanthoxeas are strewn throughout the sponge, but are most abundant near the surface. Small irregular particles that appear to be sand grains are also abundantly incorporated throughout the sponge. Surface is often very silty.

#### Spicules

Tylostyles, oxeas, and acanthoxeas; sometimes centrylote microxeas.

Tylostyles: bearing a well-formed round head or subterminal bulge. Width slowly tapering to points; pooled samples 615-1268-1923 x 7-17-28 μm (n=68; average length in each sample varies from 951 to 1671 μm).

Oxeas: straight but usually bearing a distinctive kink near the center. Centrylote swelling absent on most spicules, but weakly present on or near the kink in a few; holotype 536-839-993 x 6-10-13 μm (n=36); pooled samples 233-810-1070 x 1-10-18 μm (n=189; average length in each sample varies from 603 to 924 μm).

Acanthoxeas: generally bent in the center (centrangulate) but some gently curved. Some have a centrylote bulge on or near bend but this is often absent. Spines vary in length and density, with most samples having both longer and shorter spined spicules; sample CASIZ 66860 is atypical in that its spicules are very densely spined. Holotype: 66-88-115 x 4-6-9 (n=54); pooled samples: 44-73-132 x 3-5-11 μm (n=305; average length in each sample varies from 55 to 88 μm).

Centrylote microxeas: centrangulate like acanthoxeas, and similar in length, but thinner, with centrylote bulge more common. Usually entirely unspined. Found in only 5 of 8 samples, and much less common than acanthoxeas even when present; some previous workers have noted that these are likely immature acanthoxeas (Morrow *et al*. 2019). One seen in holotype; pooled samples: 45-61-78 x 1-3-5 μm (n=18).

#### Distribution and habitat

Uncommon from Bodega Head in Northern California to Oxnard in Southern California, and from the intertidal to 16 m in depth. Of the four samples we found, three were on shallow (<16 m) rocky ridges under kelp canopy, in one case partially buried by sand; the fourth was in a tidal pool.

## Discussion

Though not formally described, this sponge has been noted by several previous authors (see below), who often noted that it was similar to the description of *Higginsia higginissima* (Dickinson 1945) from the Sea of Cortez. This claim is difficult to assess based on Dickinson’s original description, which was very limited. Thankfully, fresh *H. higginissima* were described in much more detail by Gomez *et al*. (2002). As *H. higginissima* is branching, treelike, and has much smaller styles that are not tylote, it is clearly distinct from *H. litorea;* see also (Hofknecht 1978). No other *Higginsia* are known from the Eastern Pacific, and encrusting species are now placed in the genus *Halicnemia* rather than *Higginsia*.

We are aware of three previous reports of *Halicnemia* in the Eastern Pacific. The otherwise Atlantic *H. patera* was reported as occurring in British Columbia, albeit with no supporting data published (Austin 1985). *Halicnemia patera* differs from *H. litorea* by being disk-shaped, having longer tylostyles with multiple tyles, longer oxeas with centrylote tyles, and longer acanthostyles. As noted in the *H. montereyensis* section below, the British Columbia sponges may in fact be *H. montereyensis*.

More similar to *H. litorea* is *H. diazae* (Desqeyroux-Faundez & van Soest 1997), described from the Galapagos and reported as occurring in the Mexican Tropical Pacific (Carballo *et al*. 2019). *Halicnemia diazae* is also the only previous species of *Halicnemia* described from the intertidal. These species have considerable differences as well, however, with *H. diazae* having echinating rhabdostyles (absent in *H. litorea*), longer centrylote oxeas without a central kink, longer tylostyles, and thicker encrustations. The final species known from the Pacific is the Chilean *H. papillosa*, which has small styles (absent in *H. litorea*) and longer acanthoxeas (Desqeyroux-Faundez & van Soest 1997). The remaining species are known only from the Atlantic and Mediterranean, mostly from deep water, and all are distinguished from *H. litorea* by the presence of additional spicules and/or spicule dimensions. Most have longer acanthoxeas than *H. litorea*; an exception is *H. geniculata* from the Mediterranean; this species is a thin yellow encrustation with similar acanthoxeas to *H. litorea*, but has much thinner tylostyles and oxeas, and the oxeas are sinuous and lack the central kink (Bertolino *et al*. 2013).

This central kink has previously been reported by Morrow et al (2019), who noted that a minority of *H. patera* oxeas lack centrylote swelling but have a central kink; in the same paper, images of *H. gallica* oxeas show that some are both kinked and centrylote. These two species differ from *H. litorea* in other aspects of spicular architecture and spicular morphology. An undescribed *Halicnemia* from the Guyana shelf was found with oxeas that are both kinked and centrylote, but that sample is an unlikely conspecific due to range (van Soest 2017). It is interesting that the central kink is also shared by *H. montereyensis*, described below, despite considerable differences between that deep-water species and this shallow one. Future molecular work is needed to determine if the kinked oxea is convergent or homologous among these taxa.

*Halicnemia litorea* is easily distinguished from the other California *Halicnemia* described below. It is very different from *H. weltoni* in both spicular complement and skeletal architecture. It is more similar to *H. montereyensis* in spicule complement, though the distributions of spicule sizes for these species are non-overlapping for all spicule types, and the skeletal morphology is very different.

This species has been mentioned by several previous authors, who reported rare specimens from the intertidal in Northern and Central California (Bakus & Abbott 1980; Klontz 1989; Ristau 1977). Consistent with overall rarity, it was not found in the extensive intertidal collections made by de Laubenfels in the 1920s (de Laubenfels 1932) nor in the Southern California Bight surveys conducted between 1976-1978 (Bakus & Green 1987). Two sponges similar to *H. litorea* were found in a Southern California survey of water deeper than 50 m, and reported as “*?Desmoxyidae sp. A*” (Green & Bakus 1994). These samples were reported as having additional small tylostyles and larger acanthoxeas than *H. litorea*, but are otherwise a good match, so it is possible *H. litorea* also occurs in deep water.

Additional data could result in *H. litorea* being split into two or more species. The lengths and widths of all spicule types varied among samples, with averages falling roughly into two groups. The four samples from Northern and Central California grouped with a sample from Santa Rosa Island: these four samples have longer and thicker spicules than the three samples from mainland Southern California. These Southern, smaller-spiculed samples occur in waters with higher average surface temperatures and are separated from the others by the geographic barrier at Point Conception; the Southern population on Santa Rosa Island matches the Northern populations in morphology, but this outer island is bathed in colder water from the California current. This morphological separation is subtle, however, with some samples showing intermediate values for some spicule types, and is not supported by the genetic data. As explained in the phylogenetic results, samples show much more apparent genetic variation that seen for other species, but this does not associate with the morphological variation, as the sample from Central California groups with one of two samples from Southern California. This species is difficult to identify in the field as it is usually quite small and lacking in surface features.

“Higginsia sp. Klontz, 1989” in part (Lee et al. 2007)

### Material examined

Holotype: CASIZ 165841; other samples CASIZ 165838, CASIZ 165844; all samples collected at the “concrete block area”, Monterey Bay Canyon, Monterey, 378-379 m, 6/3/1994. Spicule measurements from CASIZ 165838 and 165844.

#### Etymology

Named for the collection location, Monterey Canyon.

#### Morphology

Samples form asymmetrical, sloping encrustations that rise into low mounds, 1-2 mm thick at one end, rising to 5-7 mm thick at the other. Prominent conules lie flat along the sponge surface and point towards thick end; sponge terminates in a cluster of conules forming a tuft. Color alive unknown; white/tan in ethanol. Holotype noted as exuding slime by collectors.

#### Skeleton

Acanthoxeas form a thick surface crust and a thinner crust along substrate; no acanothoxeas seen in between. Heads of tylostyles seen embedded along substrate, pointing up, but no complete tylostyles recovered in sections. Oxeas tightly bundled into cables 8-20 spicules thick which run at various angles, some parallel to surface, others slightly angled up; some terminate in surface conules.

#### Spicules

Tylostyles, oxeas, and acanthoxeas.

Tylostyles: bearing well-formed round heads, width slowly tapering to points. Uncommon, and with points usually broken, so lengths are an underestimate; 3672-4037-4223 x 55-60-63 μm (n=3).

Oxeas: often but not always with distinctive kink near the center. Kink is less distinct in *H. montereyensis* than in *H. litorea*, and many oxeas are completely straight; 1647-2177-2559 x 26-32-37 μm (n=30; average length of each sample separately: 2062, 2212 μm).

Acanthoxeas: a minority are bent in the center (centrangulate) but most are gently curved or straight. Very heavily spined except near tips; 118-167-202 x 14-19-23 μm (n=58; average length of each sample separately: 164, 173).

#### Distribution and habitat

Known only from a single collection, at the “concrete block area” in Monterey Canyon, Central California, 378-379 m depth.

## Discussion

The most distinctive thing about this sponge is its skeletal arrangement and overall morphology. The tight bundles of oxeas arise from the substrate at an acute angle, travel in various directions, and some terminate in conules and tufts. This overall morphology and skeleton is very similar to the description of *H. patera* in Ackers *et al*. (2007). However, a later revision indicates that this description instead pertains to *H. gallica*, which had been lumped with *H. patera* but was later resurrected (Morrow *et al*. 2019). A second species has this same morphology and skeleton: *H. wagini* (Morozov *et al*. 2018). It therefore seems likely that *H. gallica, H. wagini*, and *H. montereyensis* inherited this morphology from a common ancestor. DNA data is only available for *H. gallica* thus far, so this hypothesis remains to be tested.

*Halicnemia montereyensis* is distinguished from all previous *Halicnemia* by the dimensions of its spicules: all spicules are the longest yet reported for their type, and are much thicker than previous descriptions. The acanothoxeas of *H. montereyensis* are approached in length by *H. patera* and *H. wagini*, but the maximum recorded widths of acanthoxeas for all previous species vary from 2-12 μm, while *H. montereyensis* averages 19 μm with a maximum of 23 μm. Likewise, oxeas are longer and much thicker in *H. montereyensis* (maximum reported thickness 5-20 μm for all previously reported species, vs. an average 32 μm and maximum of 37 μm in *H. montereyensis*); the same pattern is true for tylostyles.

*Halicnemia montereyensis* is easily distinguished from the California species described here. Welton Lee identified these samples as “Higginsia sp. Klontz 1989”, together with some of the *H. litorea* described above, but these deep-water samples are quite distinct from the shallow water *H. litorea*. All three spicule types are much longer and thicker in *H. montereyensis* than *H. litorea* (size distributions are non-overlapping for both length and width for each spicule type). Gross morphology differs, with *H. montereyensis* forming the sloping mounds described above and *H. litorea* forming thin encrustations with no conules. Spicular architecture is different as well, with bundles of megascleres rising at right angles to the substrate in *H. litorea* and running nearly horizontally in *H. montereyensis*. The acanthoxeas of *H. montereyensis* are less centrangulate, have larger spines, and are more densely spined than other species described here −- except for one sample of *H. litorea* (CASIZ 66860) which shows more densely spined acanthoxeas reminiscent of *H. montereyensis*.

Most previously described species of *Halicnemia* have oxeas with one or more centrylote swellings. It is interesting that *H. montereyensis* and *H. litorea* share a central kink instead of the central swelling, despite otherwise seeming very different. As discussed above, this feature is apparently shared with two other *Halicnemia* (Morrow *et al*. 2019) but only one of these (*H. gallica*) shares the sloping morphology of *H. montereyensis.* There is a previous report of *H. patera* in British Columbia (Austin 1985). The report is hard to assess, as it is simply a species list with no descriptive data, but it is possible that the British Columbia samples are *H. montereyensis* due to the similarities with the morphological description previously attributed to *H. patera* (Ackers *et al*. 2007).

*Higginsia* sp. 1 (Lee *et al*. 2007a)

### Material examined

Holotype: CASIZ 108411, Cordell Bank, Marin County, 82 m depth, collected Sept. 1940 by “Swift, Smith, Hannah”. CASIZ 184380, Cordell Bank, Marin County (38.023, −123.4263), 55-61 m depth, collected 10/9/10, collectors not recorded.

#### Etymology

Named for Welton Lee, formerly of the California Academy of Sciences.

#### Morphology

The holotype is a roundish mass, about 50 x 30 mm, which could be described as thickly encrusting or massive. Sample CASIZ 184380 is flatter, 20 x 10 x 5 mm. Both are hispid, rugose and irregular, somewhat compressible; preserved color is white, live color is unknown.

#### Skeleton

A chaotic, halichondriid reticulation of thick styles without visible spongin. Oxeas and thin styles are loosely organized into bundles, some of which pierce the surface of the sponge to create hispidity. Acanthoxeas form a dense surface crust but are also common throughout the sponge. Small irregular particles that appear to be sand grains are also incorporated throughout the sponge.

#### Spicules

Thick styles, thin styles, long oxeas, and acanthoxeas.

Thick styles: not tylote. Could be characterized as rhabdostyles, as they often have a sharp bend near head end; 650-898-1221 x 27-36-46 μm (n=38; average length of each sample separately: 887, 1081μm).

Thin styles: sometimes with weak subterminal swelling but otherwise non-tylote. Straighter and less common than thick styles; 746-1186-1957 x 5-16-27 μm (n=12; average length of each sample separately: 1080, 1262 μm).

Long oxeas: often bent near center (centrangulate), but many are smoothly curved or straight. Usually, but not always, with weak centrylote swelling; 607-1041-2845 x 5-11-26 μm (n=34; average length of each sample separately: 887, 1177 μm).

Acanthoxeas: usually centrangulate, sometimes with weak centrylote swelling. Small spines cover the entire surface; 49-133-221 x 3-6-9 μm (n=51; average length of each sample separately: 121, 148 μm).

#### Distribution and habitat

Known only from a single collection from Cordell Bank National Marine Sanctuary, Marin, California, 55-82 m depth.

## Discussion

The differentiation between *Halicnemia* and *Higginsia* remains incompletely clear. Morrow (2019) states that *Halicnemia* have tylostyles while *Higginsia* have only styles. *Halicnemia weltoni* has only styles, but the thinner ones are sometimes subtylote and could be derived from tylostyles. *Halicnemia* are described as encrusting or disk-shaped while *Higginsia* are “erect, lamellate, massive, vasiform or lobate” (Hooper 2002a). While *H. weltoni* could be considered massive, it is similar to other massively encrusting *Halicnemia* and differs from most *Higginsia,* which are stalked and branching or upright and digitate. It is perhaps most similar to *Halicnemia diazae*, which has rhabdostyles and a skeleton described as “pluomse-dendritic” (Desqeyroux-Faundez & van Soest 1997). We consider *Halicnemia* to be a better fit, though it seems likely that future revision is needed to clearly define and separate these two genera.

The lack of tylostyles distinguishes this species from all described *Halicnemia* except *H. arcuata*, which has much shorter oxeas and acanthoxeas. Rhabdostyles are present in only two other *Halicnemia*, and in both cases they are much shorter and thinner *(H. diazae* 280-770 x 10-23 μm; *H. geniculata* 147-705 x 2-4 μm; vs. *H. weltoni* 650-1221 x 27-46 μm).

This new species can also be differentiated from named *Higginsia.* The only other Eastern Pacific *Higginsia*, *H. higginissima*, is branching and treelike with much smaller spicules (Gómez *et al*. 2002). Having thick, plain styles as the principal skeletal component also differentiates this species from most *Higginsia* described world-wide. Many *Higginsia* have oxeas only, and those with styles and oxeas usually have oxeas as the principal skeletal component and thin styles as peripheral components (e.g. *H. higgini*, *H. massalis*). The only species where styles comprise the primary skeleton (or where styles are present and descriptions of skeletal architecture are limited) have much smaller styles (*H. robusta* [styles 740 μm], *H. thielei* [styles 600-700 μm], *H. pulcherrima* [styles 420-520 μm]).

These samples were previously investigated by Lee et al. (2007a), who recognized them as distinct from anything else known from the region and referred to them as “*Higginsia* sp. 1 [of Lee]”. In the same publication, another sponge from the same location was referred to as “*Higginsia* sp. 2 [of Lee]”, and said to be perhaps conspecific with *Higginsia* sp. 1 but contaminated with foreign spicules. We investigated sample CASIZ 108398, the only sample known to be identified as *Higginsia* sp. 2 by Welton Lee, and found it to contained no acanthoxeas and seemed to be of a different order. Finally, it should be noted that CASIZ 184380 also contains fragments from the species *Forcepia macrostylosa*.

### Dichotomous Key to California Axinellida

1A. Contains polyactines (3 to 5 pointed spicules) … 2 1B. No polyactines … 4

2A. Polyactinces with bulbous tips: *Cyamon koltuni* (Sim & Bakus 1986) 2B. Polyactines without bulbous tips … 3

3A. Sponge with prominent laminate blades; polyactines with large spines on shortest clade: *Trikentrion helium* (Dickinson 1945)

3B. Sponge a thick encrustation or bushy mass; polyactines unspined or with small spines: *Cyamon neon* (de Laubenfels 1930)

4A. Contains acanthoxeas … 5 4B. No acanthoxeas … 7

5A. Acanthoxeas average more than 10 μm thick, long oxeas over 20 μm thick: *Halicnemia montereyensis sp. nov*.

5B. Acanthoxeas average less than 10 μm thick, long oxeas less than 20 μm thick … 6

6A. Acanthoxeas average 120 - 150 μm long, long oxeas without central kink: *Halicnemia weltoni sp. nov*.

6B. Acanthoxeas average 50 - 90 μm long, most long oxeas with central kink: *Halicnemia litorea sp. nov*

7A. Contains acanthostyles … 8

7B. No acanthostyles … 11

8A. Acanthostyles are subtylote with head asymmetrically placed to one side … 9

8B. Acanthostyles straight or bent, but not with subtylote with head asymmetrically placed to one side … 13

9A. Sponge is green alive and when preserved, styles average < 1000 μm in length: *Aulospongus viridans sp. nov*.

9B. Sponge is brown when preserved, surface extremely rugose, styles average > 1500 μm in length: *Aulospongus lajollaensis sp. nov*.

11A. Contains only styles or with occasional rare oxeas … 12

11B. Oxeas & styles both common, average lengths between 250 and 400 μm; sponge surface smooth: *Dragmacidon mexicanum* (de Laubenfels 1935)

12A. Multiple size categories of styles; shortest category is sharply bent near center: *Eurypon curvoclavus sp. nov*.

12B. One size category of styles, not sharply bent: *Endectyon hispitumulus sp. nov*.

13A. Many acanthostyles with severe (90 degree or more) bend near head, acanthostyles are subtylote, sponge is fan shaped: *Raspailia (Raspaxilla) hymani* (Dickinson 1945)

13B. Acanthostyles straight or with bend or curve less than 90 degrees, sponge not fan shaped … 14

14A. Acanthostyles with bend near center of shaft, average less than 150 μm long: *Eurypon curvoclavus sp. nov*.

14B. Acanthostyles straight, curved, or with bend bear head, averag more than 150 μm long … 15

15A. Sponge comprised of bushy fronds or low mounds that crumble apart, acanthostyles average over 185 μm long: *Endectyon (Endectyon) hyle* (de Laubenfels 1930)

15B. Sponge encrusting, acanthostyles average less than 185 μm long: *Endectyon hispitumulus sp. nov*.

## Conclusions

Biological diversity provides an endless supply of discovery and inspiration. In the past, Axiniellid sponges in California have been used to make advances in natural products chemistry (Aguilar-Martinez & Liaaen-Jensen 1974) and understanding archaeal symbiosis (Preston *et al*. 1996). The work reported here helps to delineate and organize the diversity in this order, creating a stable foundation for additional discoveries.

Reconciling the taxonomy of the species described here with their molecular phylogeny remains a daunting task. Our hope is that regional revisions like this one, which include both DNA and morphological data, provide insights that allow the community to work towards a global revision.

## Acknowledgements

We are grateful for the help and support of many people in UCSB’s Marine Science Institute and Diving & Boating Program, especially Robert Miller, Clint Nelson, Christoph Pierre, and Christian Orsini. NOAA provided R/V Tegula small boat support for diving operations, and Steve Lonhat (NOAA) and Shannon Myers (UCSC) were instrumental in facilitating collections in Central California. The Natural History Museum of Los Angeles’ DISCO program facilitated collections in Los Angeles County. Christina Piotrowski, Kathy Omura, Vanessa Delnavaz, and Charlotte Seid graciously provided access to their collections at the California Academy of Sciences, the Natural History Museum of Los Angeles, the Santa Barbara Natural History Museum, and the Scripps Institution of Oceanography, respectively.

